# Nonenzymatic loop-closing ligation generates RNA hairpins and is a template-free way to assemble functional RNAs

**DOI:** 10.1101/2021.11.19.469337

**Authors:** L.-F. Wu, Z. Liu, S. J. Roberts, M. Su, J. W. Szostak, J. D. Sutherland

**Affiliations:** MRC Laboratory of Molecular Biology, Francis Crick Avenue, Cambridge Biomedical Campus, Cambridge, CB2 0QH, UK; Howard Hughes Medical Institute, Department of Molecular Biology, and Center for Computational and Integrative Biology, Massachusetts General Hospital, Boston, Massachusetts 02114, United States; Department of Genetics, Harvard Medical School, Boston, Massachusetts 02115, United States

## Abstract

RNA hairpin loops are the predominant element of secondary structure in functional RNAs. The emergence of primordial functional RNAs, such as ribozymes that fold into complex structures that contain multiple hairpin loops, is generally thought to have been supported by template-directed ligation. However, template inhibition and RNA misfolding problems impede the emergence of function. Here we demonstrate that RNA hairpin loops can be synthesized directly from short RNA duplexes with single-stranded overhangs by nonenzymatic loop-closing ligation chemistry. We show that loop-closing ligation allows full-length functional ribozymes containing a hairpin loop to be assembled free of inhibitory template strands. This approach to the assembly of structurally complex RNAs suggests a plausible pathway for the emergence of functional RNAs before a full-length RNA copying process became available.

## Introduction

Functional RNAs such as ribozymes, riboswitches and aptamers – both naturally occurring and those selected by in vitro evolution – adopt folded structures (*1–5*). The prebiotic generation of structured RNA is thus crucial to the emergence of functional ribozymes during the origin of life. However, the compact, folded structure required for catalysis is incompatible with the demand for an unstructured RNA as a template for copying. Thus, a fragmentation strategy has previously been explored to assemble full-length RNAs, based on the rationale that copying the unstructured, constituent fragments individually would be less problematic than copying the structured, full-length. The question of the emergence of functional RNA could then divided into the synthesis/copying of the short fragments (*6–8*) followed by an assembly process leading to functional RNAs (*9–14*). Two strategies for the assembly process have been explored. Nicked duplex ligation to assemble full-length functional RNAs (*9, 10*) was first reported in 1966 (*15, 16*). The advantage of this strategy is that it allows many ways to position ligation junctions and template strands (splints) so as to assemble the unfolded form of the structured ribozyme (R to R^L^ in Figure 1A, top pathway). However, this strategy is inevitably subject to template inhibition (*7–10, 16*), thus product purification (*9*) or elaborate template design (*10*) are needed to enable RNA function. A further problem with the generation of complex structured RNAs is that the final structure is only realized during the folding stage (R^L^ to R in Figure 1A). In extant biology chaperones can guide the folding of functional RNAs (*17, 18*), but, absent chaperones, proper folding of complex RNAs is usually imperfect, with a substantial fraction of molecules becoming trapped in metastable mis-folded states (*19*). An alternative strategy, which leaves the fragments non-covalently joined, divides a full-length functional RNA into shorter pieces by interrupting the RNA chain within loops (R to R* in Figure 1A). This strategy was pioneered by Doudna et al. three decades ago (*11*). The non-covalently assembled complexes generated in this way could retain function, but the loose structure typically leads to some loss of activity and greater sensitivity to conditions such as elevated temperature (R* in Figure 1A) (*11*). This strategy has subsequently been explored for other ribozymes and for aptamers (*12–14*). We reasoned that if the nicked loops could be non-enzymatically sealed without disrupting the pre-structured assembly (R* to R in Figure 1A), it would offer a new strategy to directly assemble full-length structured RNAs, potentially circumventing template inhibition, misfolding and disassembly issues simultaneously (Figure 1A).

**Figure 1.**
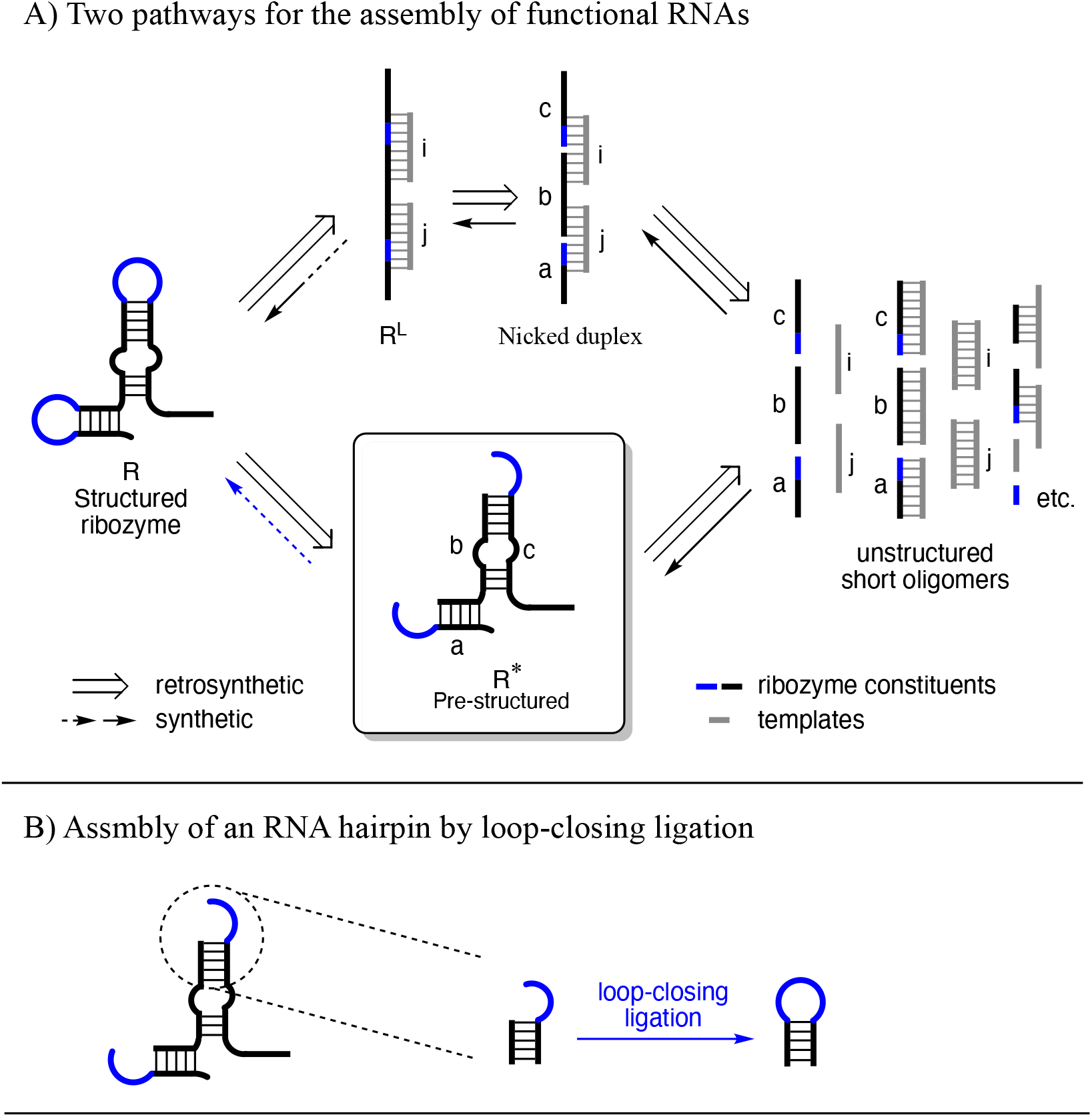
Potential loop-closing ligation constructs stem-loop hairpin structure directly. A) Conventional nicked duplex ligation strategy (top pathway) and the potential loop-closing ligation strategy (bottom pathway) to assemble full-length functional RNAs from short oligonucleotides. B) Loop-closing ligation could potentially enable a template-free way to assemble structured, functional RNAs.

We conceived of loop-closing ligation as a means of surmounting the difficulties in the assembly of active ribozymes from smaller, easier-to-replicate fragments (Figure 1B). The idea of loop-closing ligation was inspired by thinking about ligation in the context of our newly discovered nicked loop aminoacyl-transfer chemistry (*20*), as opposed to the traditional nicked duplex scenario (Figure S1). The efficiency of nonenzymatic nicked duplex RNA ligation is due to the proximity of the 3′- and activated 5′-termini of two abutting strands imparted by template binding (Figure S1-i) (*15, 16*). Similarly imparted proximity between the 3′-terminus of a primer and the activated 5′-phosphate of a monomer is the basis of template-directed RNA primer extension (*21*). In model studies of prebiotic ligation and primer extension, nucleotide activation has typically involved 5′-phosphorimidazolides (*21, 22*). These are more reactive than the triphosphates used in extant biology and their reaction with the 3′-termini of RNA strands does not require macromolecular catalysis to proceed at a reasonable rate, although this high reactivity also results in unavoidable competing background hydrolysis (*21, 22*). The proximity of the internal termini of a nicked RNA duplex can also be exploited in aminoacyl-transfer chemistry (*23*) (Figure S1-ii). Thus, if the 5′-phosphate terminus is converted into a mixed anhydride with an amino acid, aminoacyl transfer to the 3′-terminal diol occurs rather than ligation (Figure S1-ii). Remarkably, the folded-back conformation of the overhang sequence in a tRNA acceptor stem-overhang mimic also allows for interstrand aminoacyl-transfer (Figure S1-iii) (*20*) because the folded conformation of the overhang places its 3′-terminal diol in proximity to the 5′-aminoacyl-phosphate of the other strand (*20, 24*). Taken together (Figure S1-i to iii), these results suggested that this proximity might also lead to ligation if the 5′-phosphate is activated as a 5′-phosphorimidazolide (Figure 1B, Figure S1-iv). Iteration of such loop-closing ligation could allow for the assembly of complex RNA structures containing multiple stem-loops, without the need for any template to guide the assembly process.

## Results

To test the feasibility of loop-closing ligation, a 5′-phosphorimidazolide activated RNA oligonucleotide (Im-p-AGCGA-3′) together with some of the 5′-phosphorylated 5-mer (5′-p-AGCGA-3′) from which it was prepared (50 μM in total) was annealed to a 10-mer RNA (50 μM) comprising the complementary strand and a 3′-overhang (5′-<u>UCGCU</u>UGCCA-3′, complementary sequence underlined) under typical nicked duplex ligation conditions (50 mM MgCl_2_, 200 mM NaCl, 50 mM HEPES, pH 8.0) (15). The reaction mixture was incubated at 20 °C and monitored by HPLC with 260 nm UV detection. After 7 days, all the Im-p-AGCGA-3′ was consumed and a new peak appeared on HPLC chromatograms. The new peak had the same retention time as a synthetic standard of the expected 15-mer product of loop-closing ligation product. The apparent observed yield of 9 % based on total pentanucleotide corresponds to a corrected yield, based upon the percentage of Im-p-AGCGA-3′ in the 5′-phosphorimidazolide RNA preparation, of 16 %. The slow reaction rate (*t*_1/2_ = 40 h, *t*_1/2_ defined here as the “reaction half-life”, is the combined rate of first-order consumption of Im-p-AGCGA resulting from both loop-closing ligation and competing hydrolysis. See Table 1 and Table S1-S4) of the loop-closing ligation is consistent with previous reports of slow nicked duplex ligation using 5′-phosphorimidazolide RNAs (*25*). To increase the reaction rate, *N*-methylimidazole (*N*-MeIm), a nucleophilic catalyst (for detailed mechanism see Figure S2) (*26*), was added to the above reaction and found to boost the reaction rate in a concentration dependent manner. Ligation yields remained almost constant (around 30 % corrected) as *N*-MeIm concentrations were varied from 10 to 100 mM, but the reaction half-life decreased from 40 hours to 0.6 hour as the concentration of *N*-MeIm changed from 0 mM to 200 mM (Table S1). After systematically exploring other parameters including temperature, pH, concentration of MgCl_2_ and concentration of NaCl that affect the model reaction (Table S1-S4), a standard condition for loop-closing ligation (50 mM *N*-MeIm, 50 mM MgCl_2_, 200 mM NaCl and 50 mM HEPES, pH 8.0 at 20 °C) was chosen, in which the reactions proceed to completion in less than 10 hours.

**Table 1.** Model reactions of loop-closing ligation: reaction efficiency depending on the overhang sequence length and sequence identity. a) Phosphate donor sequence is Im-p-AGCGA. b) Corrected yield = Observed yield divided by the initial fraction of Im-p-AGCGA present in the pre-synthesized mixture of Im-p-AGCGA & p-AGCGA (for synthetic methods see the SI). c) Reaction half-life, *t*_1/2_, is the combined rate of first order consumption of Im-p-AGCGA resulting from both loop-closing ligation and competing hydrolysis. All yields and half-lives are average values from at least two independent experiments.

| Entry | Phosphate acceptor<sup>a</sup> | | Corrected yield<sup>b</sup> | Reaction half-life<sup>c</sup> ($T_{1/2}$, h) |
| --- | --- | --- | --- | --- |
|  | Overhang | Stem |  |  |
| 1 | UGCCA-3' | 5'-UCGCU | 30 % | 1.8 |
| 2 | CA-3' |  | 0 % | 1.4 |
| 3 | CCA-3' |  | 2 % | 1.4 |
| 4 | UCA-3' |  | 5 % | 1.5 |
| 5 | UCCA-3' |  | 15 % | 1.6 |
| 6 | ACCA-3' |  | 5 % | 1.5 |
| 7 | UGCCCA-3' |  | 13 % | 1.8 |
| 8 | UACCA-3' |  | 19 % | 1.7 |
| 9 | UCCCA-3' |  | 21 % | 1.5 |
| 10 | UUCCA-3' |  | 30 % | 2.0 |
| 11 | UGCCC-3' |  | 4 % | 1.4 |
| 12 | UGCCU-3' |  | 2 % | 1.4 |
| 13 | UGCCG-3' |  | 30 % | 1.3 |
| 14 | UGCCA(2'd)-3' |  | 2 % | 1.9 |
| 15 | UGCCA(3'd)-3' |  | 1 % | 1.6 |
| 16 | UGCCA-3' | 5'-UCGCC | 7 % | 1.5 |

Considering the variety of hairpin loops that are present in functional RNAs (27–30), we wished to explore the scope of the loop-closing ligation. Using the overhang UGCCA as a reference sequence (30 % corrected yield, Entry 1, Table 1), we investigated the effect of varying the 3′-overhang sequence. Shortening the overhang length to 2 or 3 nucleotides resulted in very poor ligation yields (< 5 % corrected, Entries 1 to 4, Table 1, Figure S3-S6). This is presumably because a 2- or 3-nucleotide long overhang cannot easily adopt a productive folded-back conformation (*27*). In accordance with the structural observations of Puglisi et al. (*24*), a UCCA overhang gave a significantly higher yield than an ACCA overhang (15 % vs. 5 % corrected, Entries 5 and 6, Table 1, Figure S7-S8). Lengthening the overhang to 6-nucleotides decreased the corrected yield to 13 % (Entry 1 versus Entry 7, Table 1, Figure S9). Changing the G at the second position of the original 5-nucleotide overhang to each of the other three nucleobases had no major effect on ligation yield or rate (Entries 8 - 10, Table 1, Figure S10-S12). However, changing the 3′-terminal A into C or U decreased the yield significantly (4 % and 2 % corrected, respectively, Entries 11 and 12, Table 1, Figure S13-S14), while changing the A into G was well tolerated (30 % corrected yield, Entry 13, Table 1, Figure S15). Changing the 3′-terminal ribonucleoside into either a 2′- or a 3′-deoxyribonucleoside suppressed the loop-closing ligation severely (< 2 % corrected yield in either case, Entries 14 and 15, Table 1, Figure S16-S17). These results show that the efficiency of the loop-closing ligation is highly dependent on the length and sequence of the overhang and the nature of the sugar moiety of the 3′-terminal nucleoside. Although a more complete data set will be required to establish potential rules for the efficiency of loop-closing ligation, it seems that both the first and last nucleotides of the overhang play significant roles, with the former likely being important in allowing a U-turn conformation and the latter in stacking to the last base pair of the stem. The effect of the sugar moiety of the 3′-terminal nucleoside might simply be explained by the lower pKa of the diol of a ribonucleoside relative to the single alcohol of a deoxynucleoside. However, it is noteworthy that the phosphodiester bonds formed by the loop-closing ligation are predominantly 2′,5′-linked rather than the canonical 3′,5′-linkage as determined by comparison to synthetic 2′,5′-linked and 3′,5′-linked standards on RNA PAGE analysis (Figure S18). The introduction of 2′,5′-linkages into functional RNAs has previously been demonstrated to be well tolerated (*31*) and will also be addressed further below.

Having optimized conditions for loop-closing ligation and partly established its scope in terms of overhang length and sequence, we decided to apply loop-closing ligation to the construction of functional RNAs, beginning with the tRNA anticodon loop. We have previously proposed that the proximity of the 5′-phosphate and 2′,3′-diol termini in a nicked loop might be responsible for both the non-enzymatic aminoacylation of the acceptor stem-overhang of a tRNA molecule and the closure of the distal anticodon loop by ligation (20). Taking the UGCCA overhang sequence we adopted in our previous aminoacylation study as a starting point, we realised that if we changed the A:U stem-closing base-pair into an A:C mismatch, the potential product of loop-closing ligation would mirror a typical tRNA anticodon loop (hepta-loop sequence CUGCCAA, Entry 16, Table 1). Indeed, when an A:C mismatch was introduced at the junction of the stem and overhang in our experimental system, loop-closing ligation was still observed (7 % corrected yield, Entry 16, Table 1, Figure S19). Interestingly, this hepta-loop product functionally resembles a tRNA anticodon loop, in that binding of this conventionally synthesized anticodon stem loop, with either a 2′,5′- or a 3′,5′-linkage between positions corresponding to residues 37 and 38 of tRNA, to oligonucleotides incorporating the complementary codon was experimentally verified by ITC (*32*) (see Methods and Figure S20). Moreover, the manner in which the loop was closed is also reminiscent of the enzymatic pathway of pre-tRNA processing after intron excision (*33*) (Figure S21).

Extending the idea of forming tRNA-like molecules from shorter oligonucleotides, we targeted a tRNA minihelix – a truncated tRNA molecule consisting of the tRNA acceptor stem-overhang and the TΨC stem-loop, and previously shown to be recognized and enzymatically aminoacylated by an aminoacyl-tRNA synthetase (*34*). Conceptually this target results from excision of the anticodon and D-stem-loops from tRNA followed by joining of the resultant termini by nicked duplex ligation (Figure 2A). Applying the latter ligation and a TΨC loop-closing ligation retrosynthetically, the tRNA minihelix can be disconnected into three fragments (Figure 2A). The RNA fragment destined to become the 5′-terminus of the tRNA minihelix (Frg-1) was 5′-FAM-labelled to enable convenient monitoring of the assembly process by gel electrophoresis. Both fragment 2 (p-Frg-2) and fragment 3 (p-Frg-3) were converted into phosphorimidazolides, Im-p-Frg-2 and Im-p-Frg-3 respectively, before mixing with Frg-1 (Figure 2B). Three FAM-labelled products (P1, P2 and P3 in observed yields of 5 %, 6 % and 0.3 % respectively) were observed after incubating all three fragments together (each at 50 uM concentration) under standard loop-closing ligation conditions (Figure 2B, Lanes 1&2 in Figure 2C). P1 represents the off-target loop-closing ligation product of Frg-1 and Im-p-Frg-3, which results from the five base-pair duplex between Im-p-Frg-2 and Im-p-Frg-3. The identity of P1 was confirmed by the fact that it is the only product formed when Frg-1 and Im-p-Frg-3 are incubated together (6 % observed yield, Lane 4&5 in Figure 2C). P2 represents the on-pathway product of nicked duplex ligation between Frg-1 and Im-p-Frg2; it is the only product observed (7 % observed yield, Lane 6&7 in Figure 2C) when Frg-1 and Im-p-Frg-2 are incubated in the presence of unactivated p-Frg-3. The third product, P3 is the expected tRNA minihelix, and this was confirmed by comparison to a standard prepared by conventional synthesis (Lane 3 in Figure 2C). A fourth product (P4) lacking a FAM label (Figure S22) was also expected from the loop-closing ligation reaction of p-Frg-2 and Im-p-Frg-3 (or reaction from Im-p-Frg-2 and Im-p-Frg-3 followed by hydrolysis of the Frg-2 phosphorimidazolide). Indeed, P4 was observed when Sybr-Gold staining was used to image the RNA gel (Figure S22) and was further verified by incubating Im-p-Frg-2 and Im-p-Frg-3 in the absence of FAM-Frg-1. These results demonstrate the power of the loop-closing ligation strategy in constructing RNA structures from short oligonucleotides. Three short RNA pieces alone were able to give four products, including the desired tRNA minihelix (P3) and two on-pathway intermediate products (P2 and P4). The formation of the off-target product (P1) in the full reaction (Figure 2B, Lanes 1&2 in Figure 2C) indicates the surprising robustness of the loop-closing ligation even in the presence of the complement strand to the overhang sequence. In a prebiotic scenario, a pool of activated short random sequence oligonucleotides could therefore have given rise to longer RNAs with structural elements more complex than simple duplexes. However, although the formation of such structures is a prerequisite for RNA function, it is not a guarantee thereof and therefore we further tested loop-closing ligation with regard to the assembly of functional, structured RNAs.

**Figure 2.**
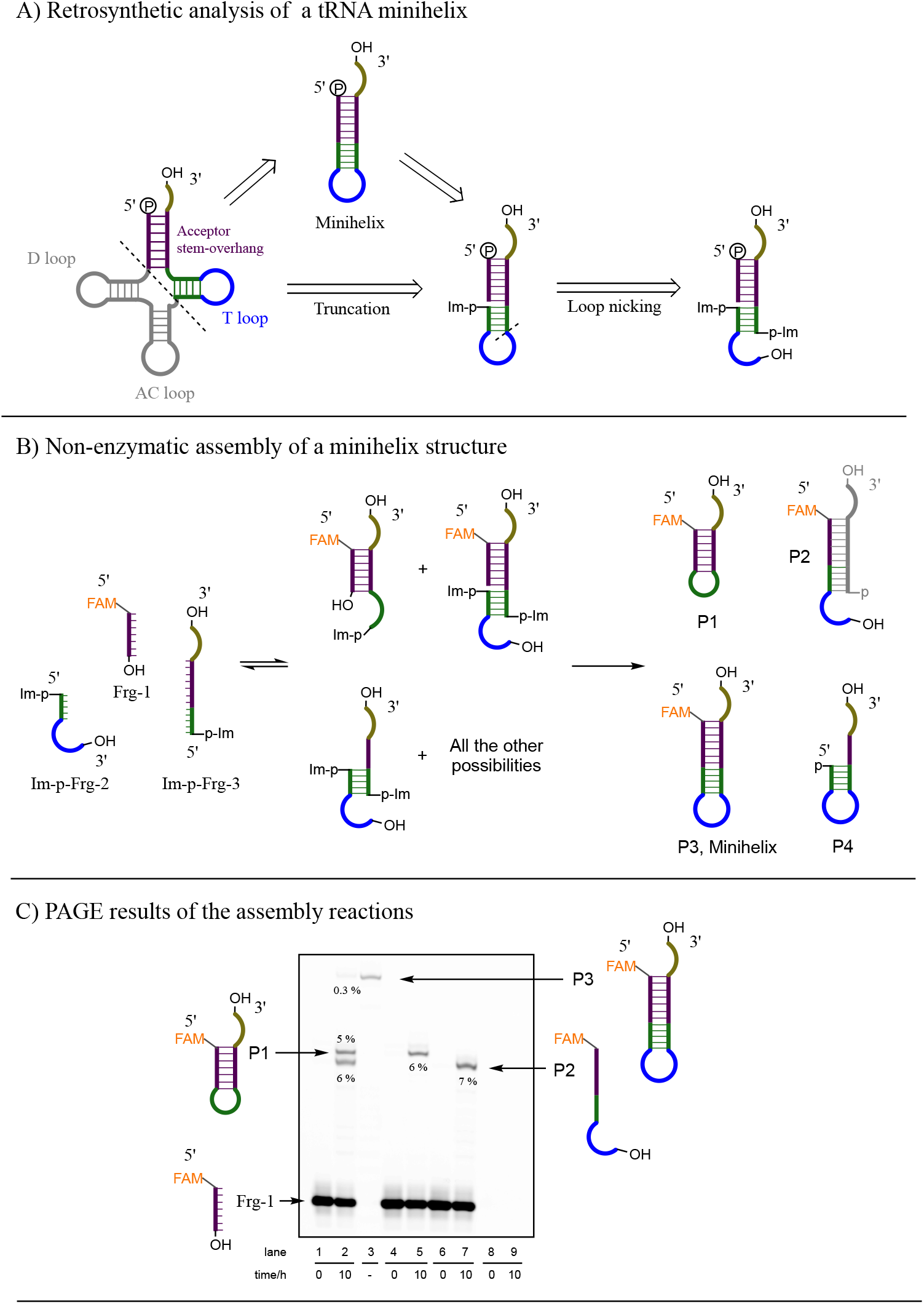
Direct assembly a tRNA minihelix by loop-closing ligation. A) The retrosynthetic truncation of tRNA and disconnection of the minihelix. B) Reaction scheme for the assembly of a tRNA minihelix. C) Representative PAGE analysis of the assembly reactions. Lane 1&2, assembly reaction of Frg-1, Im-p-Frg-2 and Im-p-Frg-3; Lane 3, authentic standard of the minihelix RNA; Lane 4&5, reaction of Frg-1 and Im-p-Frg-3; Lane 6&7, reaction of Frg-1, Im-p-Frg-2 and unactivated p-Frg-3; Lane 8&9, reaction of p-Frg-2 and Im-p-Frg-3 (no FAM-labelling oligos in this reaction). Yields are average values observed from duplicates.

In proof of principle experiments, we targeted the well-studied hammerhead ribozyme first, the catalytic core of which includes a typical hairpin stem-loop (*35*). The full-length hammerhead ribozyme was retrosynthetically disconnected in the loop region, generating a 5′-half fragment (HH-5′-Frg) and a 3′-half fragment (p-HH-3′-Frg) (Figure 3A). The HH-5′-Frg strand was labelled with a FAM fluorophore, and the p-HH-3′-Frg strand was converted into the phosphorimidazolide activated form (Im-p-HH-3′-Frg) before mixing with the HH-5′-Frag (each at 50 μM). Under our standard loop-closing ligation conditions, a 12 % yield of the full-length hammerhead ribozyme (HH-Full) was observed after 10 hours (Figure 2B, and Figure S). We then diluted the loop-closing ligation mixture 500-fold, such that the final concentration of full-length ribozyme, HH-Full, was about 12 nM, into a solution containing 1 μM of FAM-labelled hammerhead ribozyme substrate (HH-Sub). When we incubated the resulting mixture at 37 °C, the yield of the cleavage product (HH-Pdt) reached 5 % after 30 min and 16% after 90 min. As a control for the effect of loop closing ligation, a sample was prepared in which the HH-5′-Frag was mixed with unactivated p-HH-3′-Frg under the same conditions. In this control experiment we observed no visible substrate cleavage after a 90 min incubation. When the loop-closing reaction mixture was diluted by only 125-fold, the yield of the cleavage product (HH-Pdt) reached 65 % after 90 min because of the higher concentration of the ligated ribozyme (about 48 nM). However, in this case the control experiment with a 125-fold dilution of unligated HH fragments (final concentration of each fragment 400 nM) did show some enzymatic activity with 10 % substrate cleavage after 90 min, consistent with previous observations that a functional hammerhead ribozyme can be partially reconstituted by noncovalent association of fragments at high concentrations (*11–14*). We also assembled a ligase ribozyme (*36*) using a single loop-closing ligation reaction, and efficient ribozyme activity was observed directly without product purification (Figure S22). These two cases clearly demonstrate the effectiveness of the loop-closing ligation in constructing functional, full-length ribozymes in a template-free manner.

**Figure 3.**
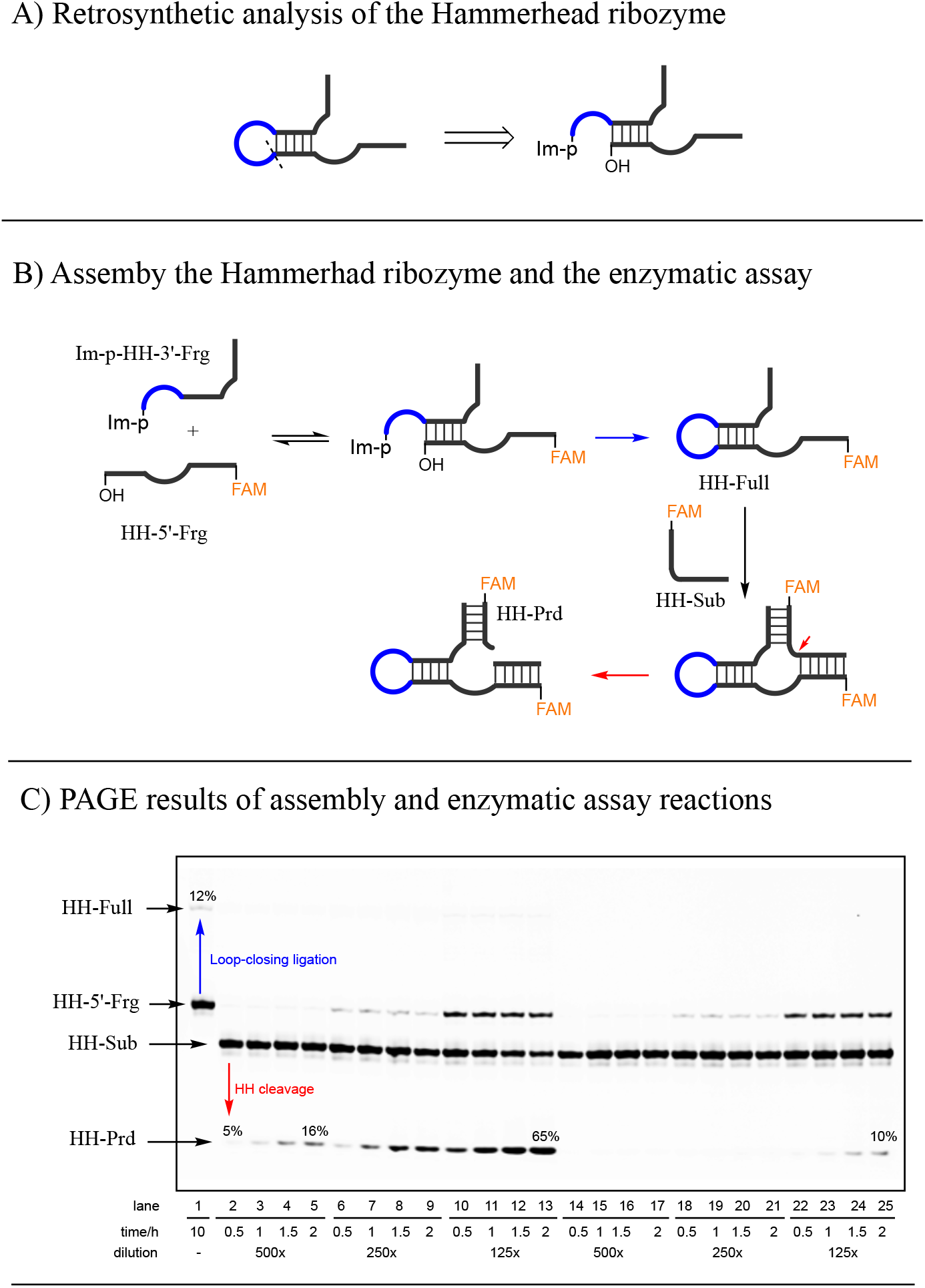
Direct assembly of the hammerhead ribozyme by the loop-closing ligation. A) The Hammerhead ribozyme was disconnected at the loop region retrosynthetically. B) Reaction scheme of the loop-closing ligation and the subsequent enzymatic cleavage of the Hammerhead substrate. C) Representative PAGE gel electrophoresis for the assembly reaction and the enzymatic assay. Lane 1, the loop-closing ligation after incubating Im-p-HH-3′-Frg (50 μM) and HH-5′-Frg (50 μM) together for 10 hours at 24 oC; Lanes 2-13, cleavage of HH-Sub by the reaction mixture after dilution; Lane 14-25, cleavage of the HH-sub by the non-covalently assembled but unligated HH fragments (without loop-closing ligation) after dilution. Yields are average values from duplicate reactions.

## Discussion

The well-known tRNA molecule has three hairpin substructures, and nearly 70 % of the nucleotides in the 16s RNA are involved in hairpins (*28, 29*). Our results suggest that hairpin loop structures could have originated directly by the loop-closing ligation of short RNA oligonucleotides. The conceptual switch from nicked duplex ligation to loop-closing ligation has two immediate consequences. Firstly, the need for external templates to join short oligonucleotides together into ribozymes is avoided (Figure 1-3), as all the strands involved become incorporated into the product by self-templating. This minimizes the problem of template oligonucleotides inhibiting product function. Secondly, the loop-closing concept makes RNA secondary structure the primary criterion to be deployed in retrosynthetic analysis of a functional RNA. This approach exploits the structural features of the target RNAs during the ligation process, thus decreases the reliance on efficient post-synthetic folding of the full length, single stranded RNA. We suggest that the shortcut of accessing RNA structures directly by loop-closing ligation could not only circumvented the template inhibition issue, but also have played key roles in the de novo emergence of structured RNAs at the origin of life. Notably, it has been demonstrated experimentally that compared to a fully random pool of RNAs, partially structured or compact, structured pools are superior sources of functional RNAs in *in vitro* selection experiments (*37, 38*). This further strengthens the potential advantage of having direct access to structured, single stranded RNAs for the emergence of function. RNA bulges and internal loops are also prevalent secondary structures (*1, 5, 39-41*), and we are currently exploring whether they too can be produced by ligation in unpaired regions of prestructured RNAs. Although we have not considered the origin of the initial pool of short oligonucleotides in this work, both untemplated synthesis and subsequent non-enzymatic templated processes could have contributed (*6–8, 21*).

Turning to specific results, the hammerhead ribozyme and a ligase ribozyme were successfully assembled from their fragments and demonstrated to be functional in situ after the loop-closing ligation. Although our data (Table 1, Figure S) suggests that the loop-closing ligation likely generated 2′,5′-linkages (3′,5′-linkages cannot be ruled out in constructs different from those in Table 1), the observation of efficient ribozyme activity (Figure 3 and Figure S22) suggested that 2′,5′-linkages in the loop regions, if present, has no dramatic impact on these two ribozymes (*31*). The relatively low yields of loop closing ligation in our studies (∼ 10 % observed yield on average) are due in part to the inefficiency of phosphoimidazolide activation, and in part to the competing hydrolysis reaction. We suggest that compatible, *in situ* activation chemistry that could maintain sets of oligonucleotides in an activated state (*22,42-43*) might drive these ligation reactions to better yields and potentially enable iterative loop-closing ligations. It is also possible that certain overhang sequences may result in higher yields of loop-closing ligation, as suggested by the varying yields in the small set of sequences examined in Table 1. The identification of such sequences will be important both for understanding the assembly of ribozymes from prebiotically available random sequence oligonucleotides, as well as for the design and assembly of ribozymes from an engineering point of view.

The non-covalent assembly of fragments into partly functional ensembles can now be seen as an evolutionary precursor of full-length ribozymes produced by loop-closing ligation of the fragments within such ensembles (*11, 14, 44-45*). We suggest that loosely structured assemblies of short oligonucleotides with diverse, but low-level function would have been accessible via dynamic assembly/disassembly of short RNA oligonucleotides at equilibrium. These loosely structured assemblies could have been pulled out-of-equilibrium and trapped in more stable structures by loop-closing ligation, thus increasing their functional activity and rendering them more robust (Figure 3C). We suggest the straightforward and economical strategy of assembling RNA structures and functions by loop-closing ligation could have been a plausible mechanism for the emergence of functions before a robust process for the replication of long RNAs was available.

## Acknowledgments

Author contributions: Loop-closing ligation was conceptualized by LFW and JDS, and was established with the assistance of ZL (2’,5’ vs 3’,5’-regioselectivity experiment), SJR (ITC experiment) and MS (part of the solid phase RNA synthesis) in the JDS group; the ribozyme work was designed by LFW and JWS, and was done in the JWS group; LFW, ZL, SJR, MS, JWS and JDS analyzed the data; LFW, JWS and JDS wrote the manuscript; the authors thank Dr. Lijun Zhou for her generous gift of the 5’-triphosphorylated RNA substrate for the ligase ribozyme, and Dr. Saurja DasGupta for his kind help with the ribozyme work. Funding: Medical Research Council (MC_UP_A024_1009), the Simons Foundation (290362 to JDS and 290363 to JWS) and National Science Foundation grant CHE-1607034 to JWS. JWS is an Investigator of the Howard Hughes Medical Institute. Competing interests: The authors declare no competing financial interests. Data and materials availability: All available data are included in the supplementary materials.

## Supporting information

### Materials and General

Reagents and solvents were obtained from *Acros Organics*, *Alfa Aesar*, *Santa Cruz Biotechnology*, *Sigma-Aldrich*, *SYNTHON Chemicals GmbH & Co. KG* and *VWR International*, and were used without further purification unless otherwise stated. For solid phase RNA synthesis, primer Support 5G for A, G, C, U or 2’-dA (with loading ∼300 μmol/g) was purchased from GE Healthcare. 3’-dA-CPG (with loading 50 μmol/g, item number 20-2004-01) was purchased from Glen Research. Phosphoramidites for RNA synthesis were purchased from Sigma-Aldrich or Link Technologies. RNA oligomers used in this study were synthesized using an ÄKTA™ oligopilot™ plus 10 (*GE Healthcare*) on a 5 to 50 μmol scale or were synthesised using an Expedite 8909 on 1 μmol scale. A *MettlerToledo* SevenEasy pH Meter S20 combined with a *ThermoFisher Scientific* Orion 8103BN Ross semi-micro pH electrode was used to measure and adjust the pH to the desired value. High-Pressure Liquid Chromatography (HPLC) was run on Dionex Ultimate 3000 (Thermo Scientific) using an Atlantis^TM^ T3, 5 μm, 4.6 x 250 mm column or an Atlantis^TM^ T3, 3 μm, 4.6 x 150 mm column. Polyacrylamide gel electrophoresis: 12 % polyacrylamide, 8 M urea gels (0.75 mm thick, 20 cm long) were run at 18 W in TBE buffer for 1 hours. FAM-labeled RNA oligomers were detected and imaged with an Amersham RGB Biomolecular Imager (GE Healthcare Life Science, Marlborough, MA) and quantified with the ImageQuant^TM^ software package (GE Healthcare Life Science, Marlborough, MA). To image unlabeled RNA oligos, the RNA gel was stained using SYBR Gold Nucleic Acid Gel Stain (Invitrogen). Oligonucleotide concentrations were determined by UV absorbance at 260 nm using a NanoDrop® ND-1000 spectrophotometer.

### Methods

#### Solid phase synthesis of RNA oligomers

After automated synthesis, RNAs were cleaved from the solid support by treating with 3 mL (for 5-50 μmol scale synthesis, if 1 μmol scale then 1.2 mL of mixture was used) of a 1:1 mixture of 28% wt NH_3_/H_2_O solution and 33% wt CH_3_NH_2_/EtOH solution at 55 °C for 30 minutes in a tube with a sealed cap. The solid was removed by filtration and washed with 50 % EtOH/H_2_O. The solution and washings were combined and evaporated to dryness under reduced pressure. Silyl protecting groups were then removed by treating the residues with 3 mL (for 5-50 μmol scale synthesis, if 1 μmol scale then 0.25 mL of mixture was used) of 1:1 mixture of triethylamine trihydrofluoride and DMSO at 55°C for 90 minutes in a tube with a sealed cap. After brief cooling at −32 °C, 30 mL of cold 50 mM NaClO_4_ in acetone was added to the solution to precipitate the RNA product. The resulting mixture was centrifuged and the pellet of RNA was re-dissolved in 10 mL of water and passed through a *Waters* Sep-Pack C18 Cartridge, 5 g sorbent (Cartridge was pre-washed with 20 mL of MeOH then 100 mL of water before sample loading, then washed with 100 mL of H_2_O, 20 mL of 10 % MeOH/H_2_O, 25 mL of 20 % MeOH/H_2_O, 25 mL of 50 % MeOH/H_2_O and 20 mL of MeOH sequentially). Eluates containing RNA were combined and lyophilized. The resulting RNA was stored as a solid or dissolved in neutral pH solution at −32 °C for future usage.

#### General procedure for chemical synthesis of Im-p-RNA

An aqueous reaction mixture (300 μL), containing the 5’-phosphoryl RNA (0.1 to 2 mM) and imidazole (50 mM), was titrated to pH 7. Then EDC (3 mg, final concentration 50 mM) was added, and the reaction mixture was incubated at room temperature. After 2 hours, 10 mL of cold 50 mM of NaClO_4_ in acetone were added to the reaction mixture to precipitate the RNA oligomers. The resulting cloudy mixture was shaken intensively and then placed in a −32 °C freezer for half an hour. The white pellet of product obtained by centrifugation was washed twice with 2 mL of cold 50 mM of NaClO_4_ in acetone, then dried in a desiccator under vacuum for half an hour. Finally, the white pellet was dissolved in 60-300 μL (making combined concentration of RNA above 0.5 mM, including both 5’-p-RNA and 5’-Im-p-RNA) of 20 mM HEPES buffer with pH 8.0 and stored at −32 °C for future usage without purification. The combined concentration of 5’-Im-p-RNA and 5’-p-RNA was measured by UV absorbance at 260 nm using NanoDrop. The yields of Im-p-RNA based on initial 5’-p-RNA ranged from 40 % to 80 % as measured by HPLC analysis.

#### Standard procedure for loop-closing ligation (Table 1, Figure S2-S17, S19)

A 50 μL reaction mixture containing Im-p-AGCGA (50 μM total, including both Im-p-AGCGA and p-AGCGA), phosphate acceptor RNA (50 μM, 5’-UCGCU<u>UGCCA</u>-3’), *N*-methylimidazole (MeIm, 50 mM, added last to initiate the catalytic reaction), NaCl (200 mM), MgCl_2_ (50 mM), cytosine (100 μM, internal reference for HPLC analysis), in HEPES buffer (50 mM, pH 8.0) was incubated at 20 °C. Aliquots (8 μL) were taken at specific time points and injected directly into an HPLC for analysis with 260 nm UV detection (Atlantis^TM^ T3, 5 μm, 4.6 x 250 mm column; flow rate 1 mL/min; LC solvents: A, 25 mM triethylammonium acetate, pH 7.5 in water and B, acetonitrile. Column compartment temperature was 25 °C). The observed yield of loop-closing ligation was calculated by comparing the above reaction to a parallel reaction run at pH 5.2 (MES buffer, 50 mM). At pH 5.2, no loop-closing ligation was observed and Im-p-AGCGA hydrolysed exclusively to p-AGCGA. Reactions with altered pH, temperature, concentration of *N-*MeIm, NaCl, MgCl_2_ were analysed similarly.

Corrected yields of the loop-closing ligation: Loop-closing ligation yields were corrected on the basis of the measured amounts of Im-p-AGCGA and 5’-p-AGCGA in the starting samples. The corrected yields represent the partition of Im-p-AGCGA into loop-closing ligation product vs. hydrolysis under a certain condition. The corrected yield does not change as the percentage of Im-p-AGCGA varies between synthetic batches, but the observed yields do.

#### Regioselectivity (2’,5’-versus 3’,5’-phosphodiester) of the loop-closing ligation in the model reactions (Figure S18)

1 μL of the above loop-closing ligation reaction was quenched with 40 μL of stop buffer (6 M urea in TEB buffer with 100 mM EDTA, pH 8.0). An all-3’,5’-linkage authentic standard and an authentic standard with one 2’,5’-linkage at the loop-closing position were prepared in stop buffer at 0.5 μM. 2 μL of the quenched and standard solutions, were analysed by PAGE. The regioselectivity of the newly formed phosphodiester bond was assigned by comparing the reactions to the standards by imaging after SYBR Gold Nucleic Acid Gel Staining.

#### Binding of the anticodon hairpin loop to its cognate codon (Figure S20)

A 17-nt anticodon loop sequence 5’-<u>GUCGC</u>CU**UCC**A*A<u>GCGAC</u>-3’ (stem sequences are underlined, the anticodon is shown in bold, and the position where the loop was closed is indicated by a star) was used. A 5-nt sequence U**GGA**A containing the corresponding GGA codon was used as a surrogate messenger RNA. Prior to use, the RNAs were refolded by heating (10 min at 95 °C) and cooling to room temperature over 1.5 hours. ITC buffer: 50 mM MgCl_2_, 1M NaCl, 50 mM HEPES, pH 7.0. ITC measurements were performed at 10 °C using a GE Healthcare MicroCal Auto-ITC-200. The concentrations of the anticodon loop titrants in the injection syringe were 404 μM for the canonical 3’,5’-linkage at A*A position of the anticodon loop, or 363 μM for the anticodon loop with a 2’,5’-linkage at the A*A position, respectively. The concentration of the 5nt sequence (U**GGA**A) in the cell was 36.4 μM. Data collected for each titration experiment were then fit to a single binding site model. *K*_d_ values were determined to be 2.5 μM for the canonical anticodon loop with a 3’,5’-linkage at the A*A position, and 9.4 μM for the anticodon loop with a 2’,5’-linkage at the A*A position (see Figure S21 for detailed parameters). This gave a good fit for both the binding of anticodon loop with a 3’,5’ linkage (N=0.79, *K*_d_ =2 .5 uM, dH = −31.2 kcal/mol and -TdS = 23.9 kcal/mol) and a 2’,5’ linkage (N=0.76, *K*_d_ = 9.4 uM, dH = −33.5 kcal/mol and -TdS = 27.0 kcal/mol) to the 5nt oligonucleotide. Oligos used for ITC were purified by a Varian Prep Star preparative HPLC system with Varian Pro Star UV/vis detector and a Water Atlantis T3 Prep OBD 5 μM 19×250 mm column. Solvent A: 20 mM pH 7 NH_4_OAc; Solvent B: Acetonitrile. 0 min 7% B, 30 min 40% B, 35 min 90% B, 40 min 90% B. Fractions containing the product were lyophilised and then checked for purity by analytical HPLC.

#### Assembly of a tRNA minihelix structure by loop-closing ligation and nicked duplex ligation (Figure 1 and Figure S22)

To a 10 μL reaction mixture containing 5’-FAM-AUUA<u>GGAGAUG</u>-3’ (FAM-Frg-1, 25 μM), 5’-Im-p-<u>GAGGG</u>UUUGAGA-3’ (Im-p-Frg-2, 25 μM in total, including 5’-Im-p-GAGGGUUUGAGA-3’ and 5’-p-GAGGGUUUGAGA-3’), 5’-Im-p-<u>CCCUUCAUCUCC</u>ACCA-3’ (Im-p-Frg-3, 25 μM in total, including 5’-Im-p-CCCUUCAUCUCCACCA-3’ and 5’-p-CCCUUCAUCUCCACCA-3’), NaCl (200 mM), MgCl_2_ (50 mM) in HEPES buffer (50 mM, pH 8.0) was added *N*-methylimidazole (MeIm, 50 mM), followed by incubation at 25 °C. Aliquots (0.5 μL) were taken at specific time points and quenched in 25 μL of stop buffer (6 M urea in TBE buffer with 100 mM EDTA, pH 8.0). 2 μL of the quenched solution was analysed by PAGE. Observed yields were quantified according to the relative amounts of FAM-labelled oligomers by gel imaging. Products without FAM-labelling was visualised, but not quantified, after SYBR Gold Nucleic Acid Gel Staining.

#### Assembly the full-length hammerhead ribozyme by the loop-closing ligation and the subsequent enzymatic assay (Figure 2)

A 10 μL reaction mixture containing 5’-FAM-ACCUGUCUGAUGA<u>GCAA</u>G-3’ (HH-5’-Frg, 50 μM), 5’-Imp-UUAU<u>CUUGC</u>GAAACCGU-3’ (Im-p-HH-3’-Frg, 50 μM, including 5’-Imp-UUAUCUUGCGAAACCGU-3’ and 5’-p-UUAUCUUGCGAAA-CCGU-3’), *N*-methylimidazole (MeIm, 50 mM, added last to initiate the catalytic reaction), NaCl (200 mM), MgCl_2_ (50 mM) in HEPES buffer (50 mM, pH 8.0) was incubated at 25 °C. A control reaction was run in parallel by replacing Im-p-HH-3’-Frg with unactivated p-HH-3’-Frg (5’-p-UUAUCUUGCGAAACCGU-3’, 50 μM). After 10 hours, 0.5 μL of the reaction solution was diluted in 20 μL of water, then 1 μL of the diluted solution was quenched in 9 uL of stop solution (6 M urea in TEB buffer with 100 mM EDTA, pH 8.0). 2 μL of the quenched solution was analysed by PAGE. Observed yields of the full-length hammerhead ribozyme (HH-Full) were quantified according to the relative amounts of FAM-labelled oligomers by gel imaging. **Enzymatic assay**: The loop-closing reaction mixture was diluted 50, 25, or 12.5 times, respectively, in water, and dilutions of the control reaction without loop-closing ligation were also prepared. Then, 1 μL of the previously diluted reaction mixtures were used to prepare 10 μL of a solution also containing 5’-FAM-AAACGGUCACAGGU-3’ (HH-Sub, 1 μM), NaCl (200 mM), MgCl_2_ (10 mM) and HEPES buffer (50 mM, pH 7.0). Each solution was incubated at 37 °C, and 1 μL of reaction mixture was quenched by adding to 9 μL of stop solution (6 M urea in TEB buffer with 100 mM EDTA, pH 8.0) at t = 0.5, 1, 1.5 and 2 hours, respectively. 2 μL of the quenched solution was analysed by PAGE. Yields of cleavage of HH-Sub (5’-FAM-AAACGGUCACAGGU-3’) to the 8-nt product (HH-Ptd, 5’-FAM-AAACGGUC>p) were quantified by gel imaging.

#### Assembly a full-length Joyce ligase ribozyme by the loop-closing ligation and the subsequent enzymatic assay (Figure S23)

A 10 μL reaction mixture containing 5’-FAM-UAAAGUUGUUAUCACU-<u>CGUAG</u>UUCCA-3’, (Lig-5’-Frg, 50 μM), 5’-Imp-<u>CUACG</u>UUAUGGAUGGGUU-GAAGUAU-3’, (Im-p-Lig-3’-Frg, 50 μM, including 5’-Imp-CUACGUUAUGGA-UGGGUUGAAGUAU-3’ and 5’-p-CUACGUUAUGGAUGGGUUGAAGUAU-3’), *N*-methylimidazole (*N*-MeIm, 50 mM, added last to initiate the catalytic reaction), NaCl (200 mM), MgCl_2_ (50 mM) in HEPES buffer (50 mM, pH 8.0) was incubated at 25 °C. A control reaction was run in parallel by replacing Im-p-Lig-3’-Frg with unactivated p-Lig-3’-Frg (5’-p-CUACGUUAUGGAUGGGUUGAAGUAU-3’, 50 μM). After 10 hours, 0.5 μL of the reaction solution was diluted in 20 μL of water, then 1 μL of the diluted solution was quenched in 9 uL of stop solution (6 M urea in TEB buffer with 100 mM EDTA, pH 8.0). 2 μL of the quenched solution was analysed by PAGE. Yields of the full-length ligase ribozyme (Lig-Full) were quantified by fluorescence gel imaging.

##### Ligase ribozyme assay

A 10 μL enzymatic reaction mixture containing ppp-GAGACCGCAACUUA (Lig-Sub, 4 μM), NaCl (200 mM), MgCl_2_ (50 mM), HEPES buffer (50 mM, pH 8.0) and ∼ 0.8 μM of Lig-Full was prepared. The Lig-Full solution (2 μL) was added last. The same procedure was applied to the control reaction without loop-closing ligation. The resulting solutions were incubated at 48 °C for 6 hours, then 1 μL of each reaction mixture was quenched by addition to 9 μL of stop solution (6 M urea in TBE buffer (I don’t think you have defined TBA buffer) with 100 mM EDTA, pH 8.0). 2 μL of the quenched solution was analysed by PAGE. The yield of the enzymatic ligation product (Lig-Ptd) was quantified by gel imaging. Products without FAM-label were visualised, but not quantified, by SYBR Gold Nucleic Acid Gel Staining

---

**Figure S1.**
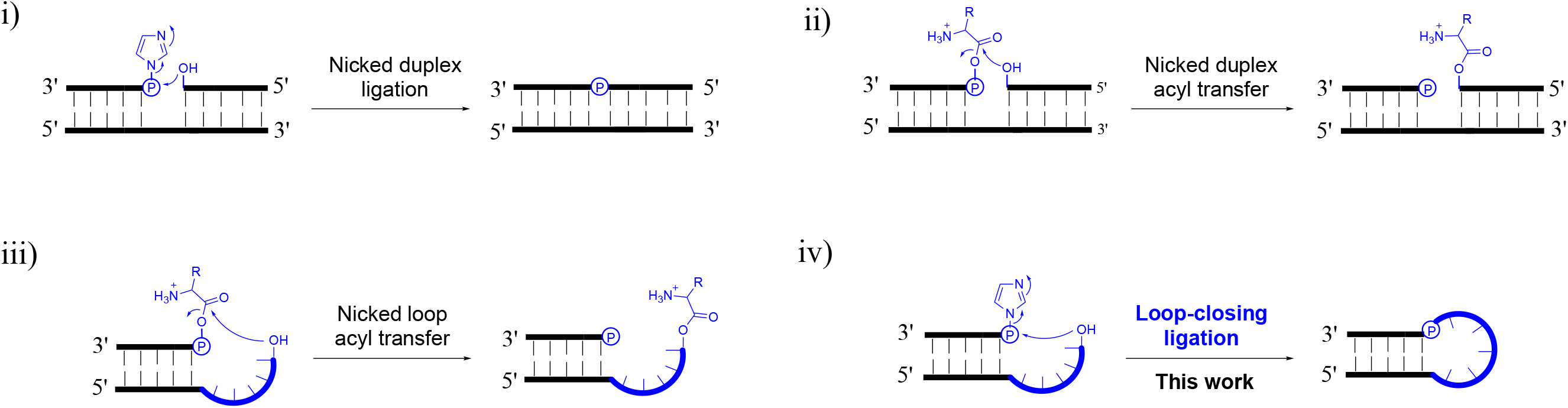
Loop-closing ligation compared to ligation and acyl-transfer reactions on a nicked duplex and a nicked loop. i) The proximity of 5’- and 3’-ends in a nicked duplex facilitates ligation using 5’-phosphorimidazolide activation, and also, ii) facilitates acyl transfer chemistry from a 5’-mixed anhydride. iii) Similarly, the proximity of 5’- and 3’-ends in a nicked loop facilitates acyl transfer chemistry from a 5’-mixed anhydride, which suggested that iv) the same proximity might facilitate ligation chemistry in a nicked loop using 5’-phosphorimidazolide activation.

**Figure S2.**
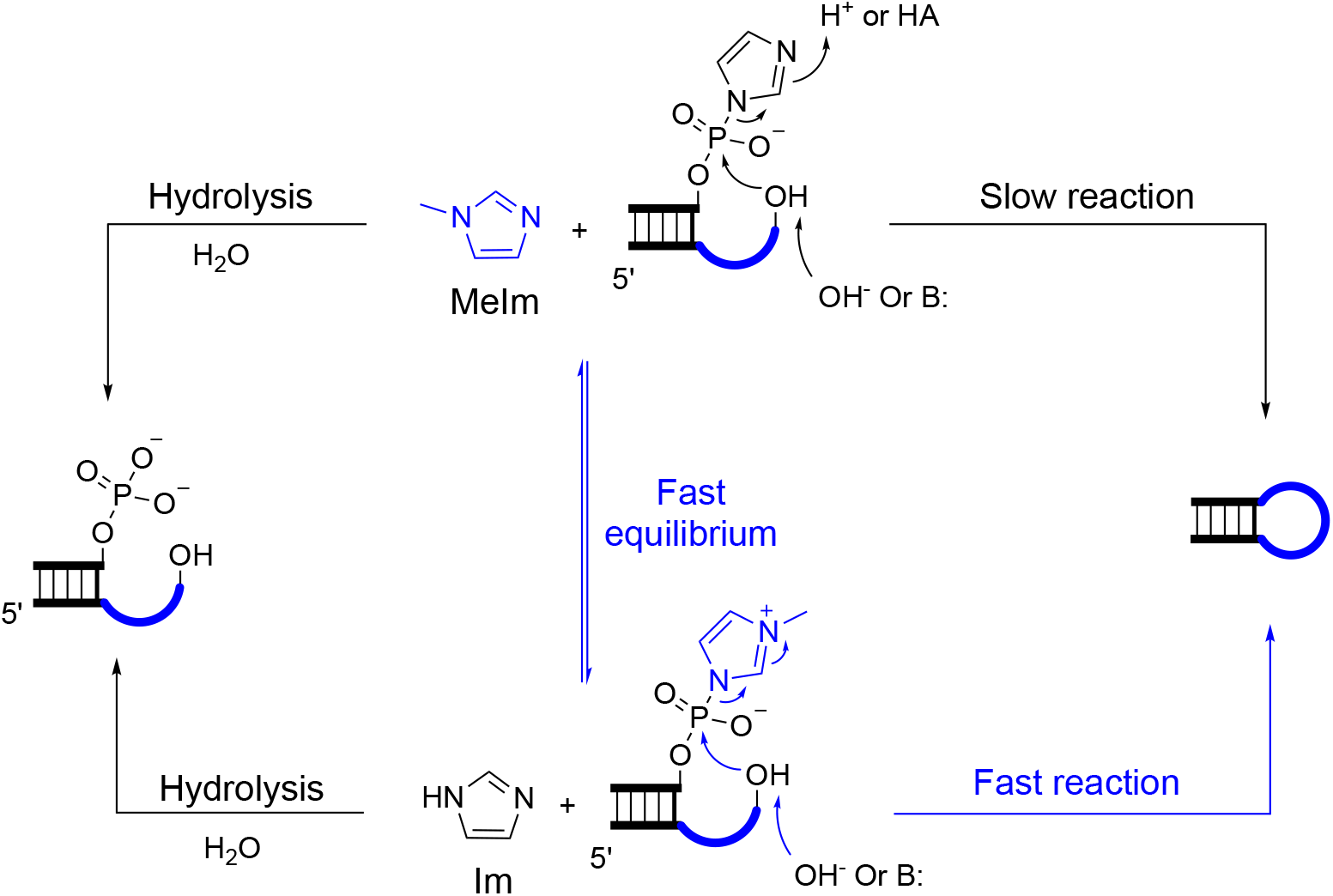
Scheme for catalysis of loop-closing ligation by *N*-methylimidazole. The equilibrium between Im-p-AGCGA + *N-*MeIm and *N-*MeIm-p-AGCGA + Im could also happen off-duplex, and all the other possible association and dissociation equilibria of RNA strands are omitted for simplicity. Standard reaction condition: 50 μM each of the phosphate donor and phosphate acceptor RNA strands, *N-*MeIm 50 mM, MgCl_2_, 50 mM, NaCl 200 mM, HEPES 50 mM, pH 8.0 at 20 °C. Im, imidazole; *N-* MeIm, *N*-methylimidazole; B:, general base; HA, general acid. The overhang sequence is highlighted in blue.

**Figure S3.**
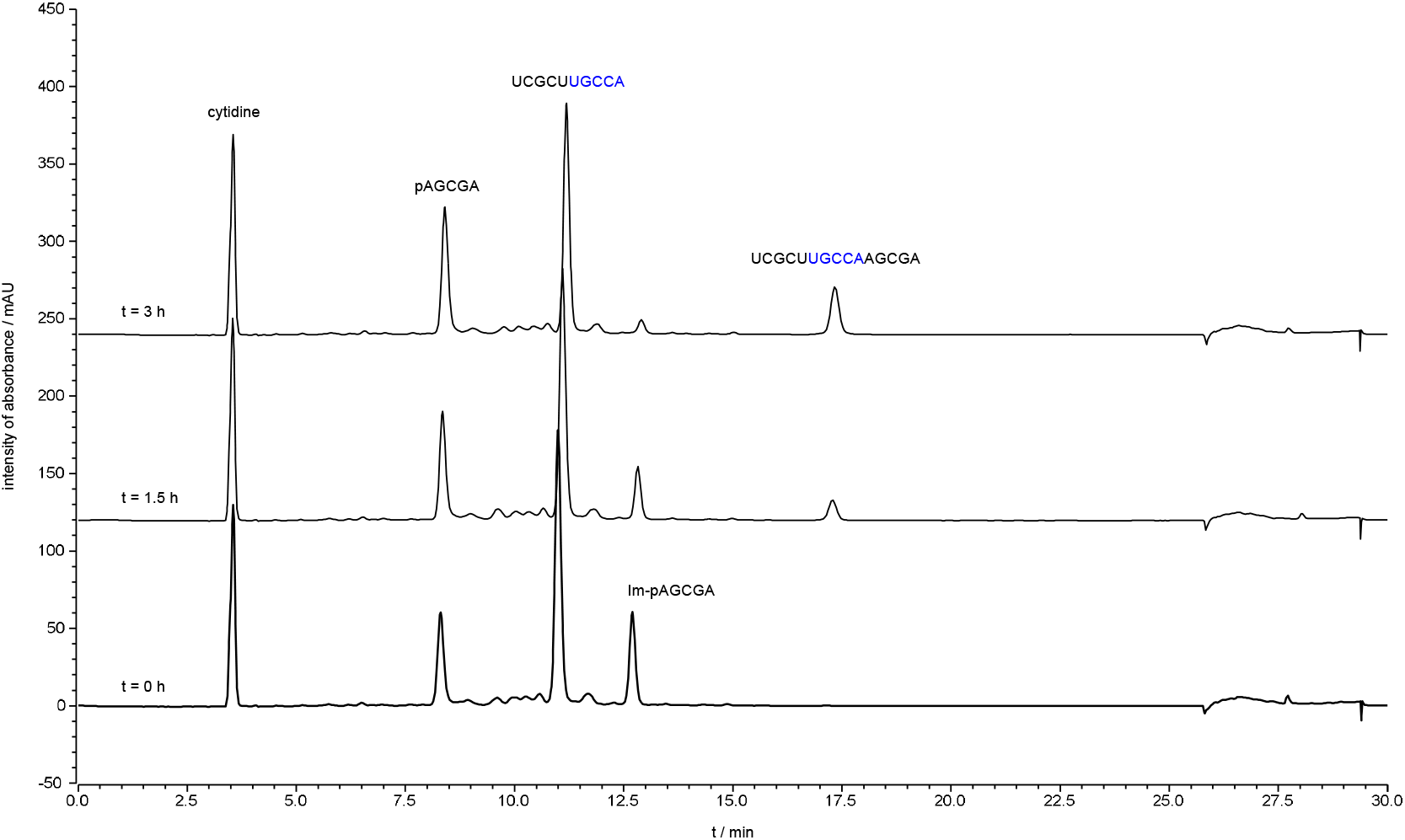
Stacked HPLC traces of loop-closing ligation with UGCCA overhang. Loop duplex sequence: `3’AGCGAp-Im` 5’UCGCUUGCCA Loop-closing ligation was monitored by HPLC with 260 nm UV detection. The solution was incubated at 20 °C and aliquots of 8 μL were injected into an HPLC at different time points. Peaks for the phosphate donor, phosphate acceptor strands and the product of loop-closing ligation are indicated. Conditions: 50 μL of reaction mixture, containing the phosphate donor strand (including Im-p-AGCGA and p-AGCGA, in total 50 μM), the phosphate acceptor strand (5’-UCGCUUGCCA-3’, 50 μM), cytidine (internal standard, 200 μM), NaCl (200 mM), MgCl_2_ (50 mM), *N-*MeIm (50 mM) and HEPES (50 mM, pH 8), was incubated at 20 °C.

**Figure S4.**
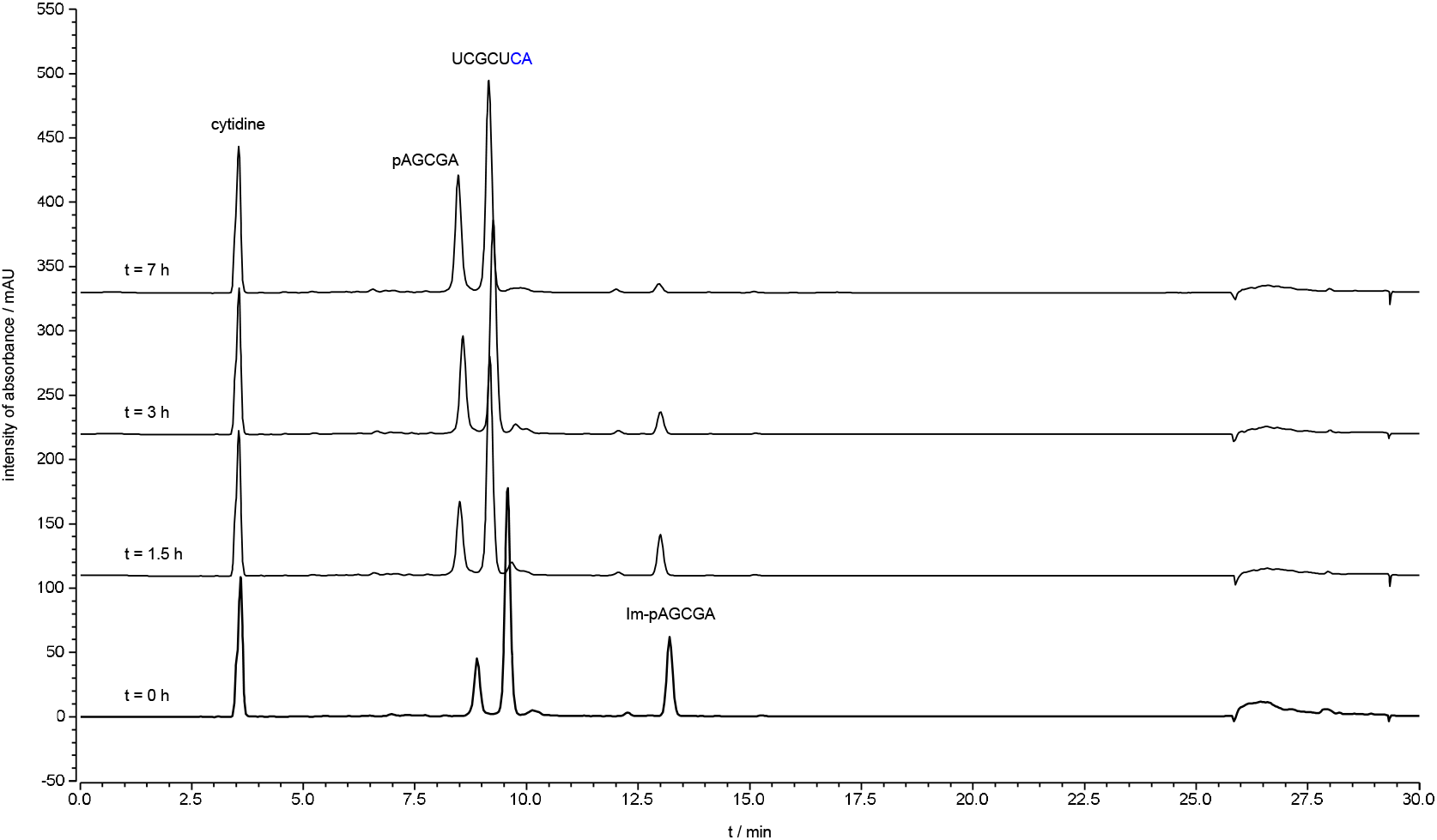
Stacked HPLC traces of loop-closing ligation with CA overhang. Loop duplex sequence: `3’AGCGAp-Im` `5’UCGCUCA` Loop-closing ligation was monitored by HPLC with 260 nm UV detection. The solution was incubated at 20 °C and aliquots of 8 μL were injected into an HPLC at different time points. Peaks for the phosphate donor, phosphate acceptor strands and the product of loop-closing ligation are indicated. Conditions: 50 μL of reaction mixture, containing the phosphate donor strand (including Im-p-AGCGA and p-AGCGA, in total 50 μM), the phosphate acceptor strand (5’-UCGCUCA-3’, 50 μM), cytidine (internal standard, 200 μM), NaCl (200 mM), MgCl_2_ (50 mM), *N-*MeIm (50 mM) and HEPES (50 mM, pH 8), was incubated at 20 °C.

**Figure S5.**
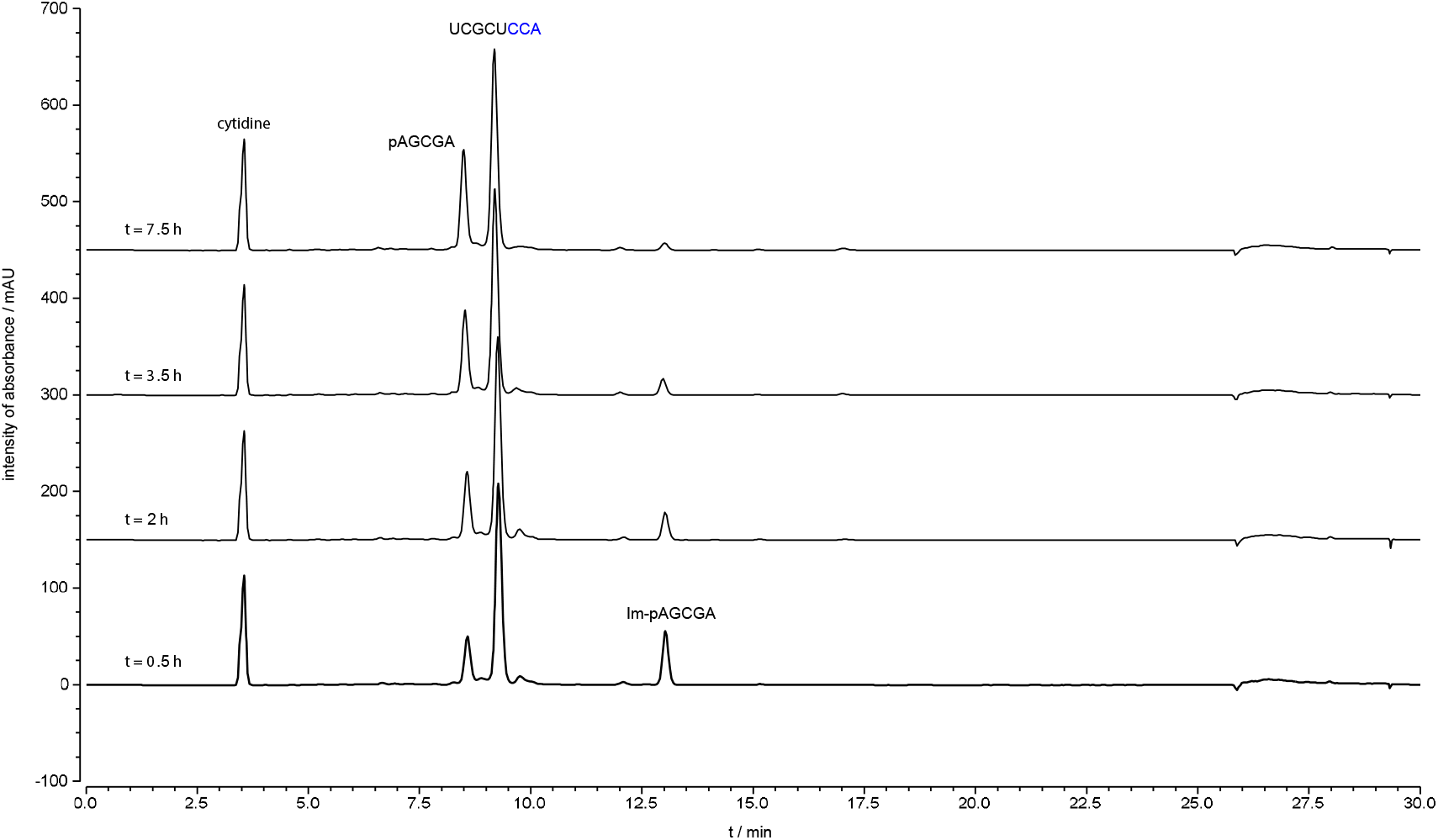
Stacked HPLC traces of loop-closing ligation with CCA overhang. Loop duplex sequence: `3’AGCGAp-Im` 5’UCGCUCCA Loop-closing ligation was monitored by HPLC with 260 nm UV detection. The solution was incubated at 20 °C and aliquots of 8 μL were injected into an HPLC at different time points. Peaks for the phosphate donor, phosphate acceptor strands and the product of loop-closing ligation are indicated. Conditions: 50 μL of reaction mixture, containing the phosphate donor strand (including Im-p-AGCGA and p-AGCGA, in total 50 μM), the phosphate acceptor strand (5’-UCGCUCCA-3’, 50 μM), cytidine (internal standard, 200 μM), NaCl (200 mM), MgCl_2_ (50 mM), *N-*MeIm (50 mM) and HEPES (50 mM, pH 8), was incubated at 20 °C.

**Figure S6.**
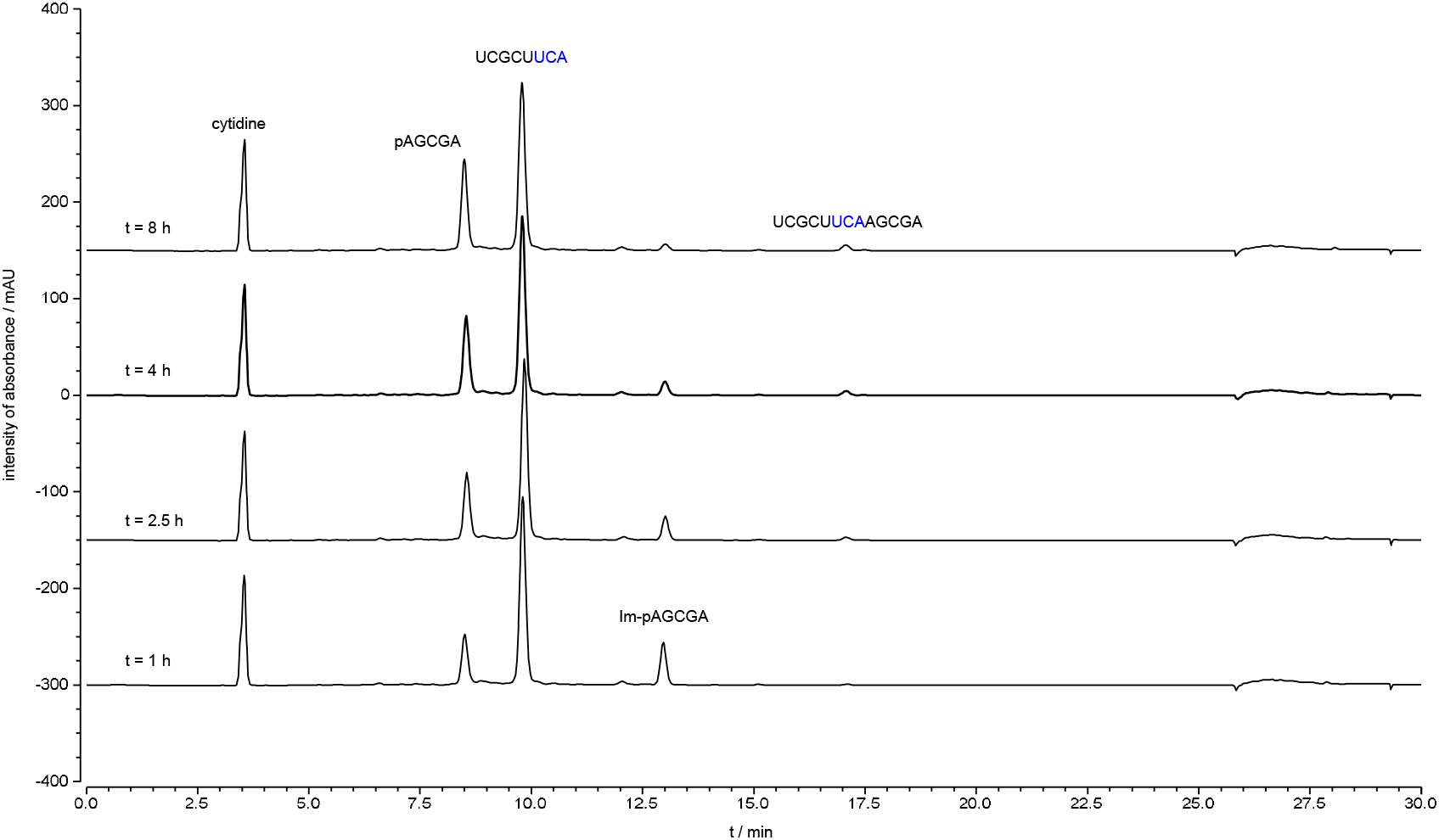
Stacked HPLC traces of loop-closing ligation with UCA overhang. Loop duplex sequence: `3’AGCGAp-Im` 5’UCGCUUCA Loop-closing ligation was monitored by HPLC with 260 nm UV detection. The solution was incubated at 20 °C and aliquots of 8 μL were injected into an HPLC at different time points. Peaks for the phosphate donor, phosphate acceptor strands and the product of loop-closing ligation are indicated. Conditions: 50 μL of reaction mixture, containing the phosphate donor strand (including Im-p-AGCGA and p-AGCGA, in total 50 μM), the phosphate acceptor strand (5’-UCGCUUCA-3’, 50 μM), cytidine (internal standard, 200 μM), NaCl (200 mM), MgCl_2_ (50 mM), *N*-MeIm (50 mM) and HEPES (50 mM, pH 8), was incubated at 20 °C.

**Figure S7.**
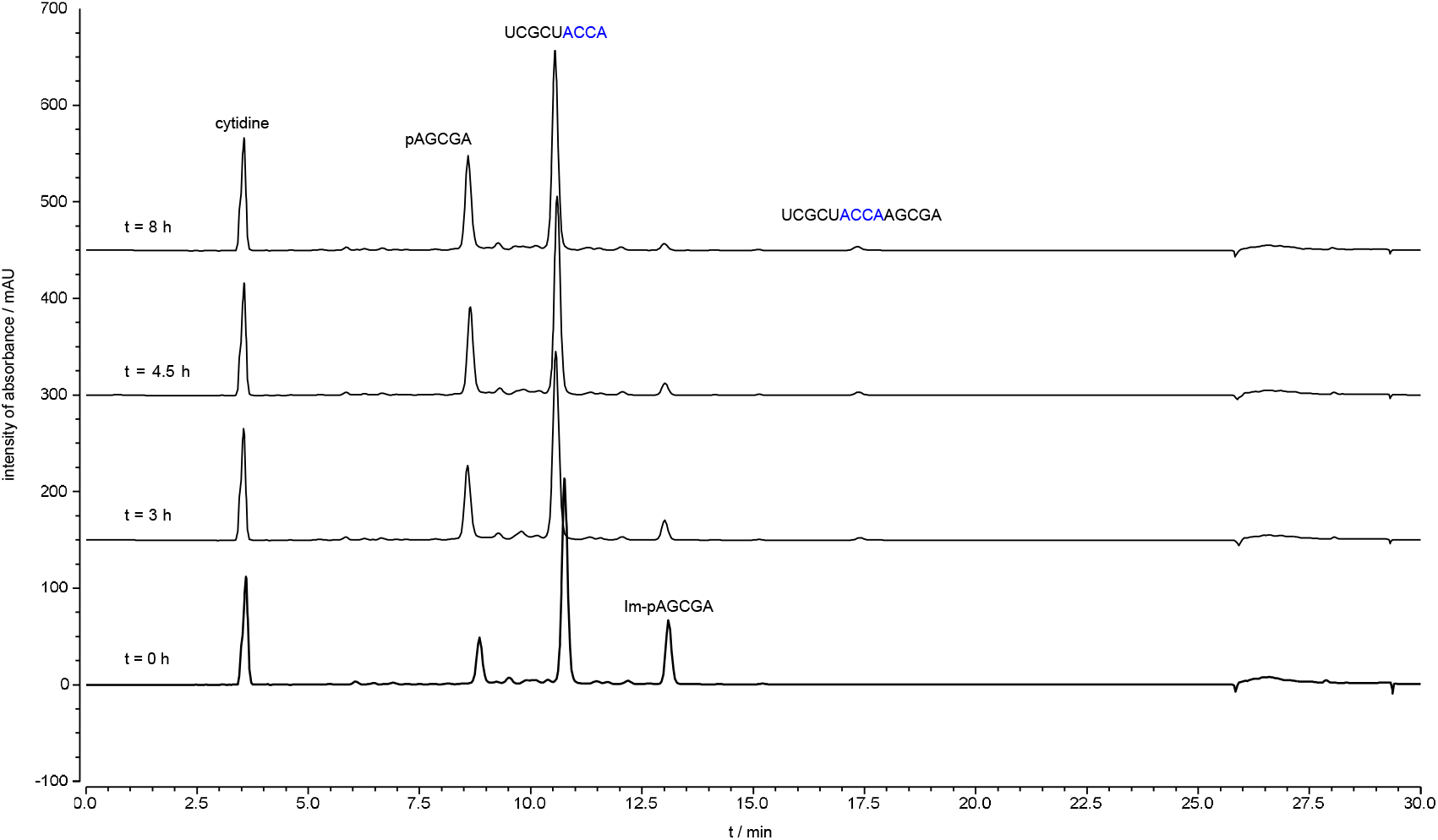
Stacked HPLC traces of loop-closing ligation with ACCA overhang. Loop duplex sequence: `3’AGCGAp-Im` 5’UCGCUACCA Loop-closing ligation was monitored by HPLC with 260 nm UV detection. The solution was incubated at 20 °C and aliquots of 8 μL were injected into an HPLC at different time points. Peaks for the phosphate donor, phosphate acceptor strands and the product of loop-closing ligation are indicated. Conditions: 50 μL of reaction mixture, containing the phosphate donor strand (including Im-p-AGCGA and p-AGCGA, in total 50 μM), the phosphate acceptor strand (5’-UCGCUACCA-3’, 50 μM), cytidine (internal standard, 200 μM), NaCl (200 mM), MgCl_2_ (50 mM), *N*-MeIm (50 mM) and HEPES (50 mM, pH 8), was incubated at 20 °C.

**Figure S8.**
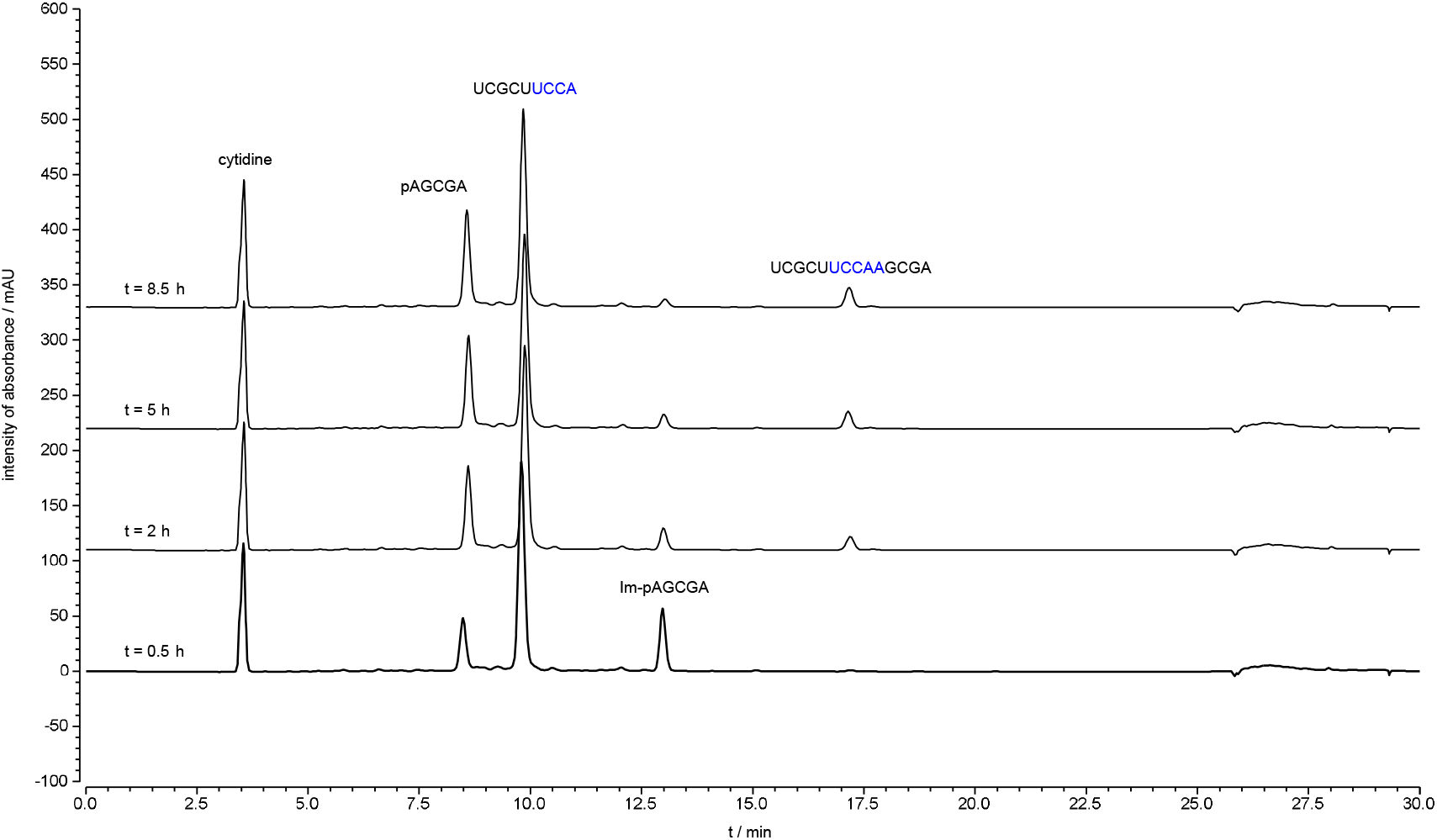
Stacked HPLC traces of loop-closing ligation with UCCA overhang. Loop duplex sequence: `3’AGCGAp-Im` 5’UCGCUUCCA Loop-closing ligation was monitored by HPLC with 260 nm UV detection. The solution was incubated at 20 °C and aliquots of 8 μL were injected into an HPLC at different time points. Peaks for the phosphate donor, phosphate acceptor strands and the product of loop-closing ligation are indicated. Conditions: 50 μL of reaction mixture, containing the phosphate donor strand (including Im-p-AGCGA and p-AGCGA, in total 50 μM), the phosphate acceptor strand (5’-UCGCUUCCA-3’, 50 μM), cytidine (internal standard, 200 μM), NaCl (200 mM), MgCl_2_ (50 mM), *N*-MeIm (50 mM) and HEPES (50 mM, pH 8), was incubated at 20 °C.

**Figure S9.**
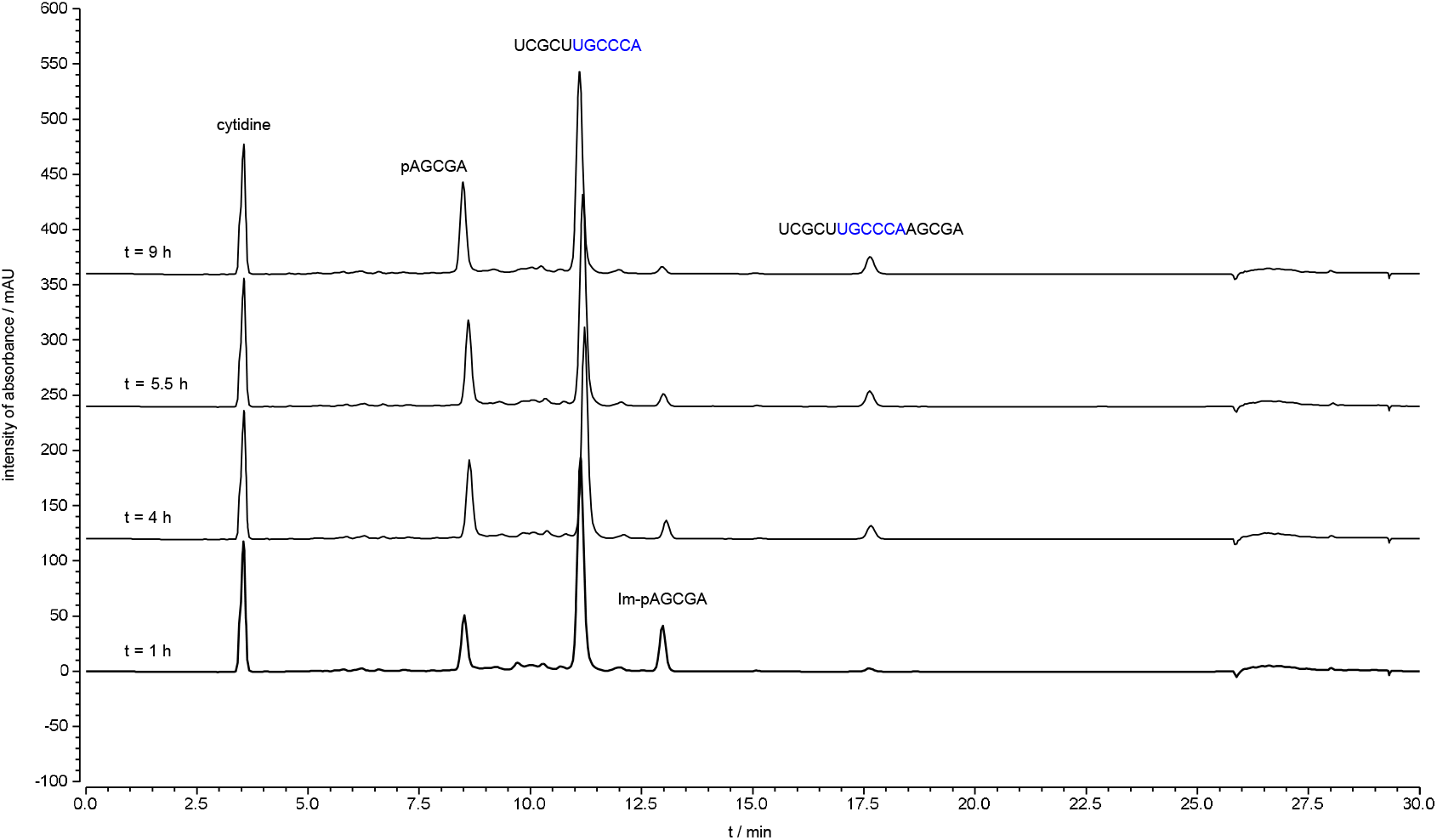
Stacked HPLC traces of loop-closing ligation with UGCCA overhang. Loop duplex sequence: `3’AGCGAp-Im` 5’UCGCUUGCCCA Loop-closing ligation was monitored by HPLC with 260 nm UV detection. The solution was incubated at 20 °C and aliquots of 8 μL were injected into an HPLC at different time points. Peaks for the phosphate donor, phosphate acceptor strands and the product of loop-closing ligation are indicated. Conditions: 50 μL of reaction mixture, containing the phosphate donor strand (including Im-p-AGCGA and p-AGCGA, in total 50 μM), the phosphate acceptor strand (5’-UCGCUUGCCCA-3’, 50 μM), cytidine (internal standard, 200 μM), NaCl (200 mM), MgCl_2_ (50 mM), *N*-MeIm (50 mM) and HEPES (50 mM, pH 8), was incubated at 20 °C.

**Figure S10.**
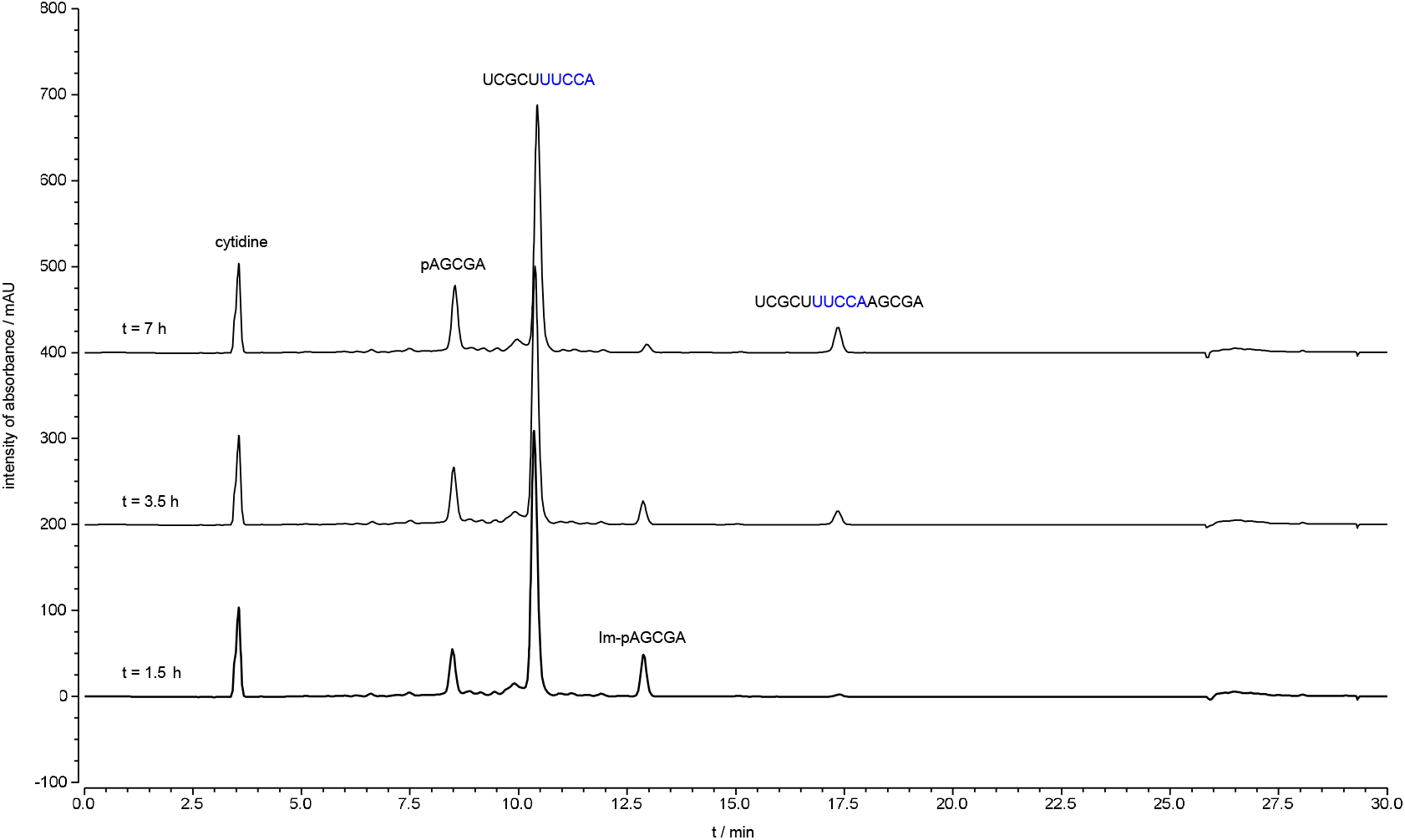
Stacked HPLC traces of loop-closing ligation with UUCCA overhang. Loop duplex sequence: `3’AGCGAp-Im` 5’UCGCUUUCCA Loop-closing ligation was monitored by HPLC with 260 nm UV detection. The solution was incubated at 20 °C and aliquots of 8 μL were injected into an HPLC at different time points. Peaks for the phosphate donor, phosphate acceptor strands and the product of loop-closing ligation are indicated. Conditions: 50 μL of reaction mixture, containing the phosphate donor strand (including Im-p-AGCGA and p-AGCGA, in total 50 μM), the phosphate acceptor strand (5’-UCGCUUUCCA-3’, 50 μM), cytidine (internal standard, 200 μM), NaCl (200 mM), MgCl_2_ (50 mM), *N*-MeIm (50 mM) and HEPES (50 mM, pH 8), was incubated at 20 °C.

**Figure S11.**
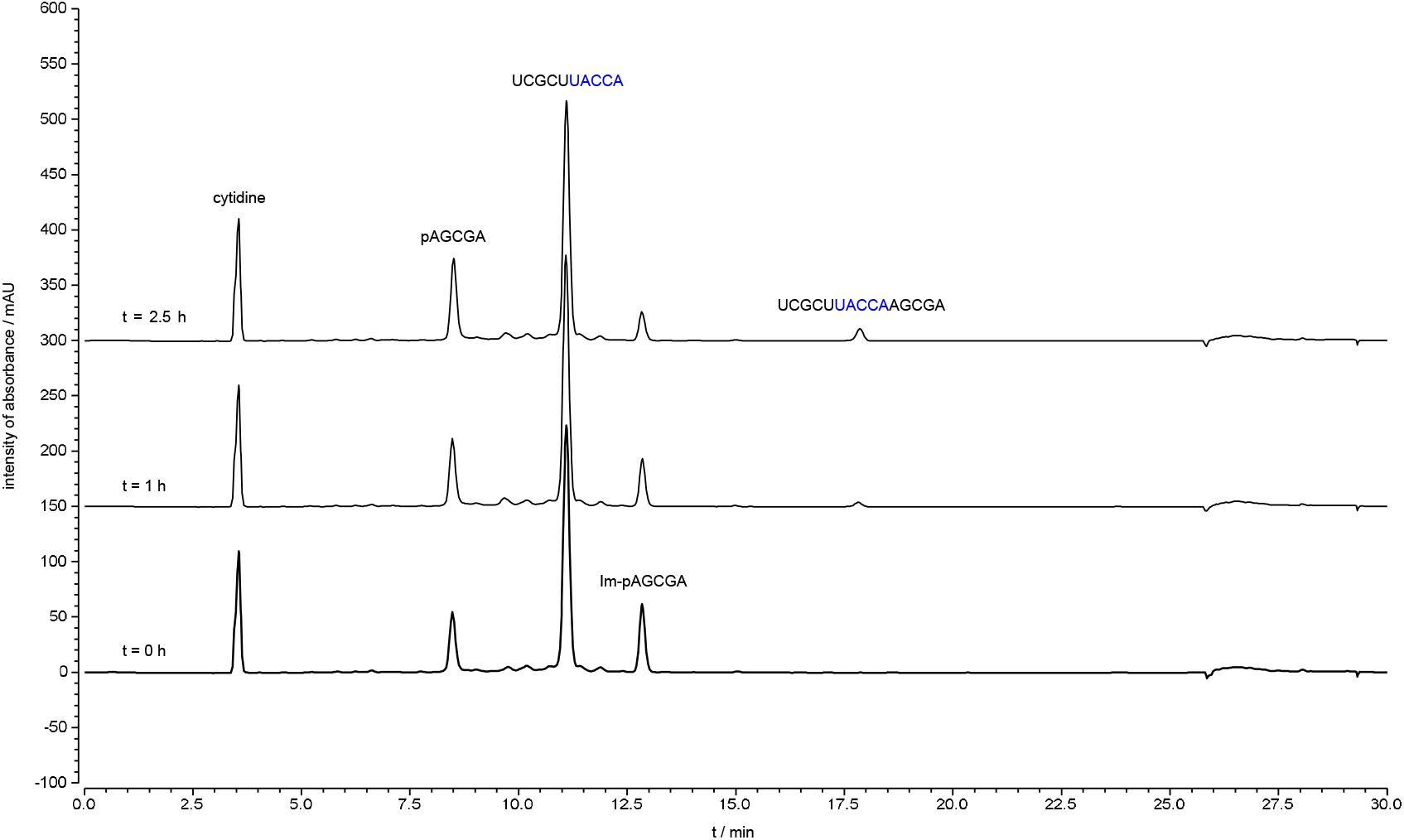
Stacked HPLC traces of loop-closing ligation with UACCA overhang. Loop duplex sequence: `3’AGCGAp-Im` 5’UCGCUUACCA Loop-closing ligation was monitored by HPLC with 260 nm UV detection. The solution was incubated at 20 °C and aliquots of 8 μL were injected into an HPLC at different time points. Peaks for the phosphate donor, phosphate acceptor strands and the product of loop-closing ligation are indicated. Conditions: 50 μL of reaction mixture, containing the phosphate donor strand (including Im-p-AGCGA and p-AGCGA, in total 50 μM), the phosphate acceptor strand (5’-UCGCUUACCA-3’, 50 μM), cytidine (internal standard, 200 μM), NaCl (200 mM), MgCl_2_ (50 mM), *N*-MeIm (50 mM) and HEPES (50 mM, pH 8), was incubated at 20 °C.

**Figure S12.**
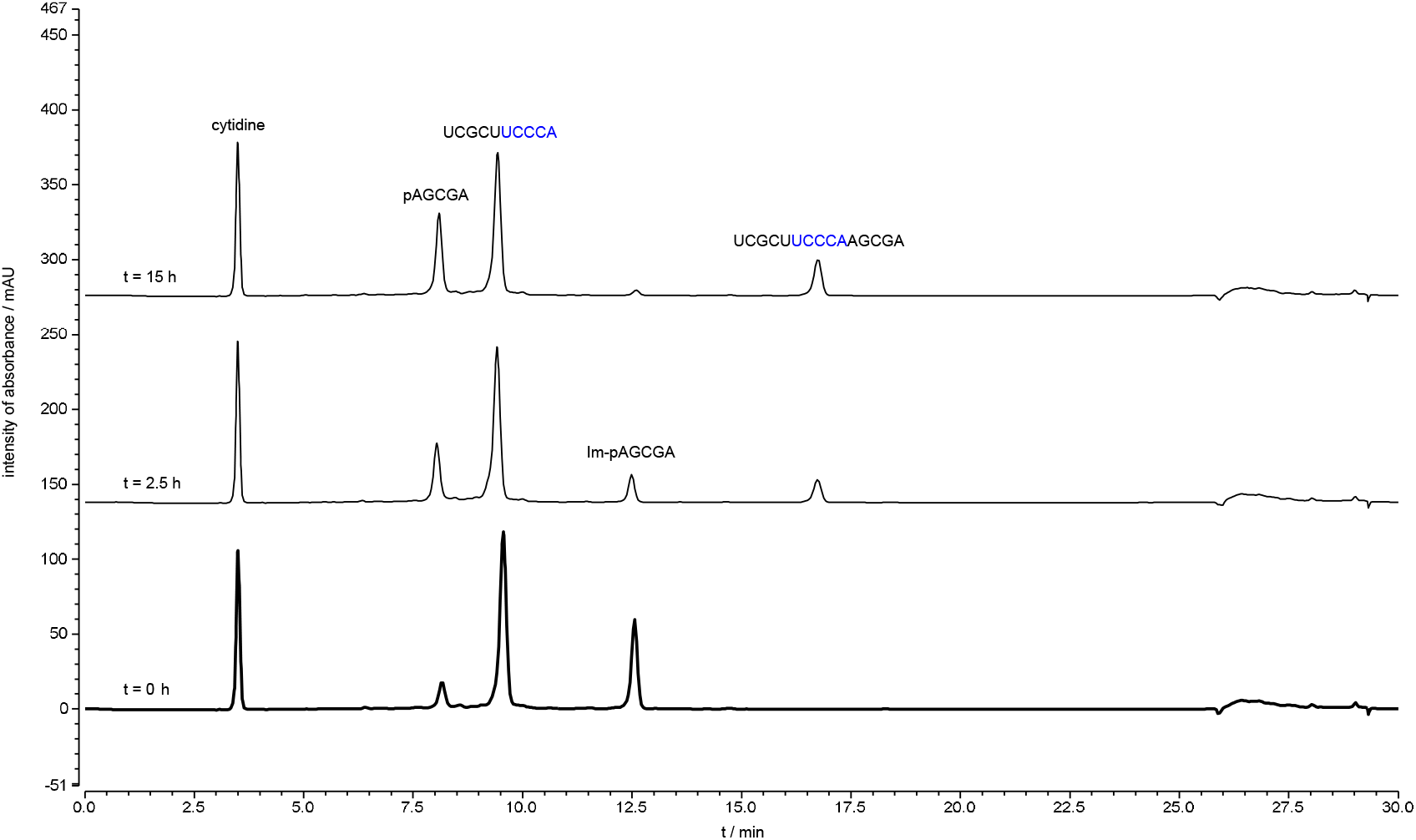
Stacked HPLC traces of loop-closing ligation with UCCCA overhang. Loop duplex sequence: `3’AGCGAp-Im` 5’UCGCUUCCCA Loop-closing ligation was monitored by HPLC with 260 nm UV detection. The solution was incubated at 20 °C and aliquots of 8 μL were injected into an HPLC at different time points. Peaks for the phosphate donor, phosphate acceptor strands and the product of loop-closing ligation are indicated. Conditions: 50 μL of reaction mixture, containing the phosphate donor strand (including Im-p-AGCGA and p-AGCGA, in total 50 μM), the phosphate acceptor strand (5’-UCGCUUCCCA-3’, 50 μM), cytidine (internal standard, 200 μM), NaCl (200 mM), MgCl_2_ (50 mM), *N*-MeIm (50 mM) and HEPES (50 mM, pH 8), was incubated at 20 °C.

**Figure S13.**
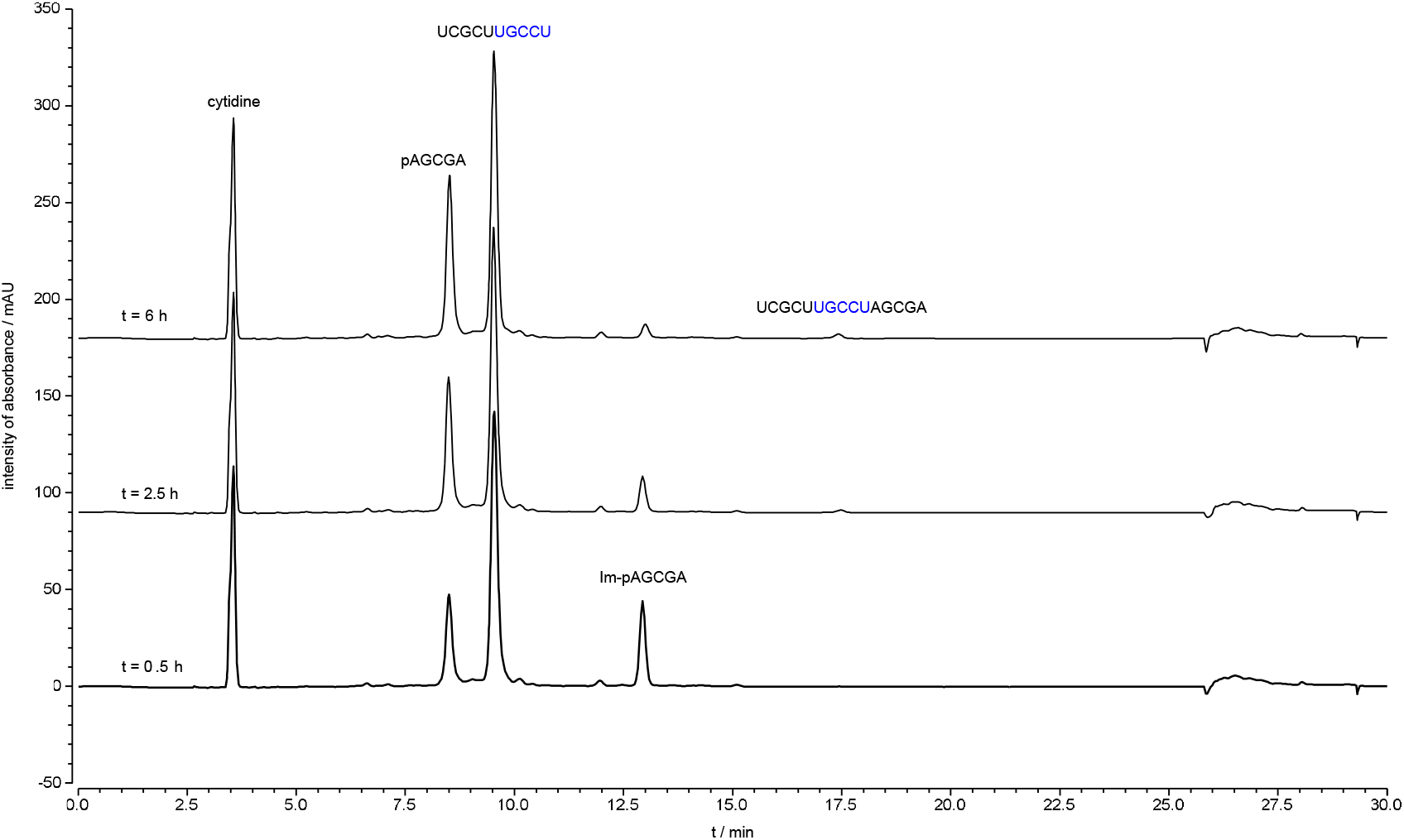
Stacked HPLC traces of loop-closing ligation with UGCCU overhang. Loop duplex sequence: `3’AGCGAp-Im` 5’UCGCUUGCCU Loop-closing ligation was monitored by HPLC with 260 nm UV detection. The solution was incubated at 20 °C and aliquots of 8 μL were injected into an HPLC at different time points. Peaks for the phosphate donor, phosphate acceptor strands and the product of loop-closing ligation are indicated. Conditions: 50 μL of reaction mixture, containing the phosphate donor strand (including Im-p-AGCGA and p-AGCGA, in total 50 μM), the phosphate acceptor strand (5’-UCGCUUGCCU-3’, 50 μM), cytidine (internal standard, 200 μM), NaCl (200 mM), MgCl_2_ (50 mM), *N*-MeIm (50 mM) and HEPES (50 mM, pH 8), was incubated at 20 °C.

**Figure S14.**
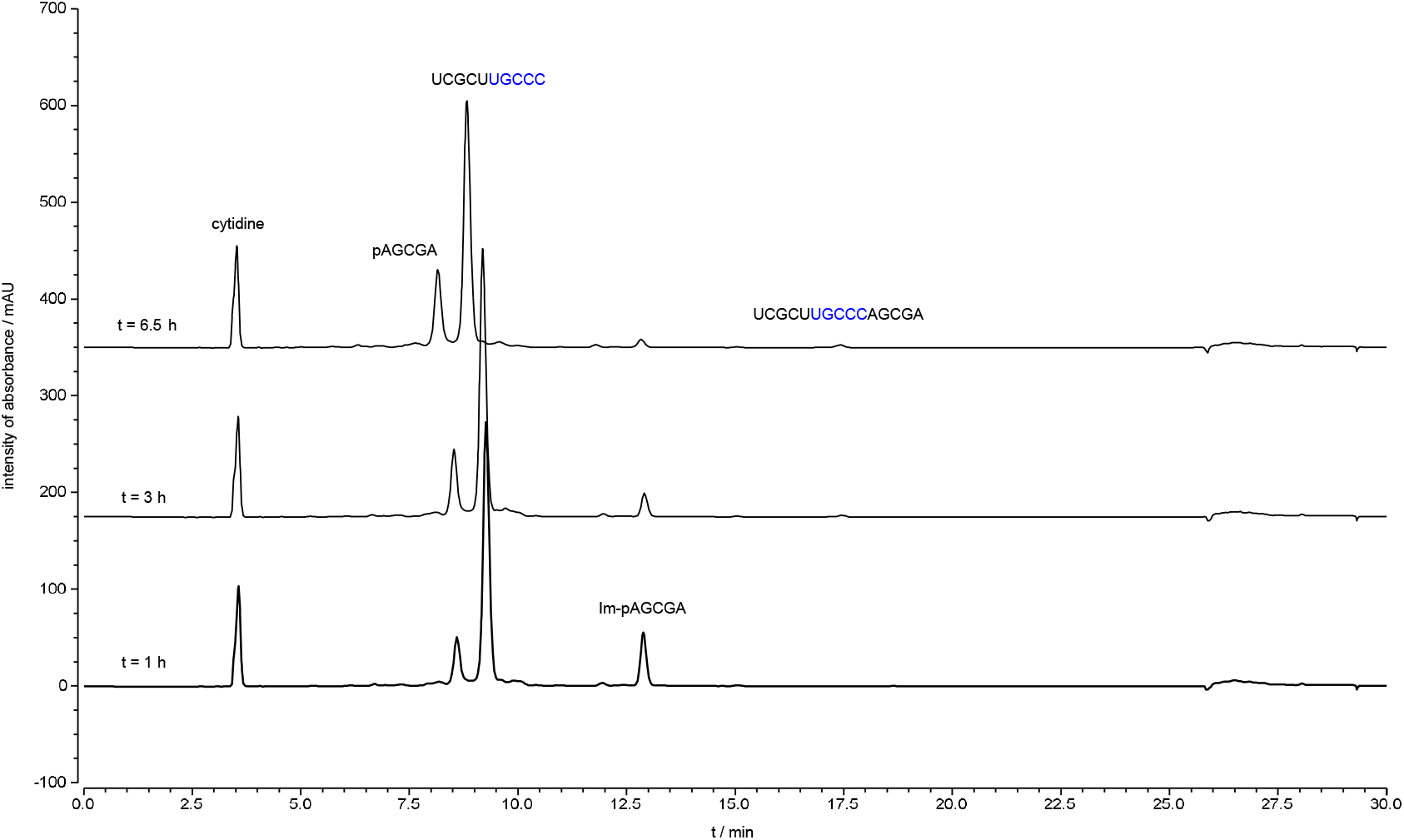
Stacked HPLC traces of loop-closing ligation with UGCCC overhang. Loop duplex sequence: `3’AGCGAp-Im` 5’UCGCUUGCCC Loop-closing ligation was monitored by HPLC with 260 nm UV detection. The solution was incubated at 20 °C and aliquots of 8 μL were injected into an HPLC at different time points. Peaks for the phosphate donor, phosphate acceptor strands and the product of loop-closing ligation are indicated. Conditions: 50 μL of reaction mixture, containing the phosphate donor strand (including Im-p-AGCGA and p-AGCGA, in total 50 μM), the phosphate acceptor strand (5’-UCGCUUGCCC-3’, 50 μM), cytidine (internal standard, 200 μM), NaCl (200 mM), MgCl_2_ (50 mM), *N*-MeIm (50 mM) and HEPES (50 mM, pH 8), was incubated at 20 °C.

**Figure S15.**
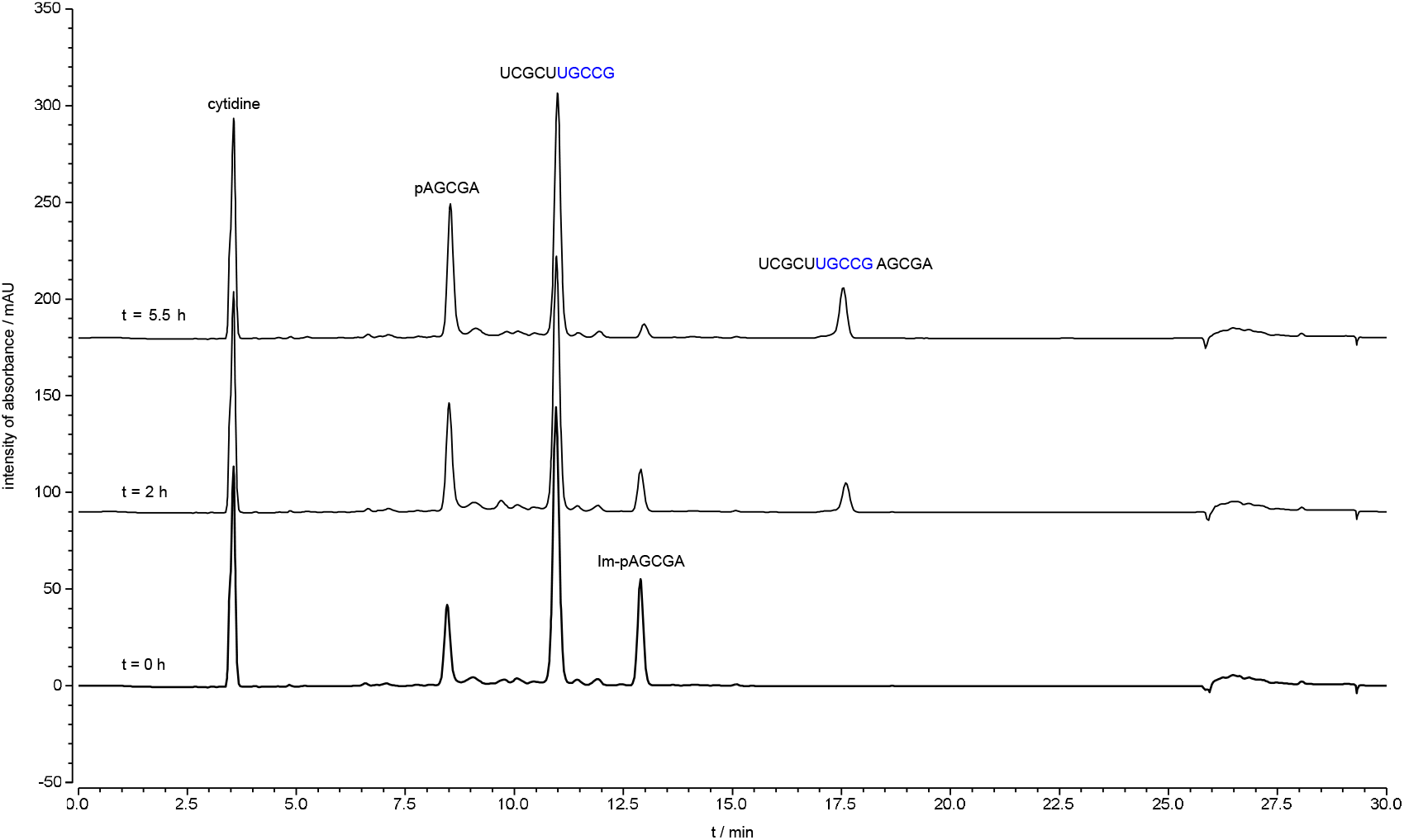
Stacked HPLC traces of loop-closing ligation with UGCCG overhang. Loop duplex sequence: `3’AGCGAp-Im` 5’UCGCUUGCCG Loop-closing ligation was monitored by HPLC with 260 nm UV detection. The solution was incubated at 20 °C and aliquots of 8 μL were injected into an HPLC at different time points. Peaks for the phosphate donor, phosphate acceptor strands and the product of loop-closing ligation are indicated. Conditions: 50 μL of reaction mixture, containing the phosphate donor strand (including Im-p-AGCGA and p-AGCGA, in total 50 μM), the phosphate acceptor strand (5’-UCGCUUGCCG-3’, 50 μM), cytidine (internal standard, 200 μM), NaCl (200 mM), MgCl_2_ (50 mM), *N*-MeIm (50 mM) and HEPES (50 mM, pH 8), was incubated at 20 °C.

**Figure S16.**
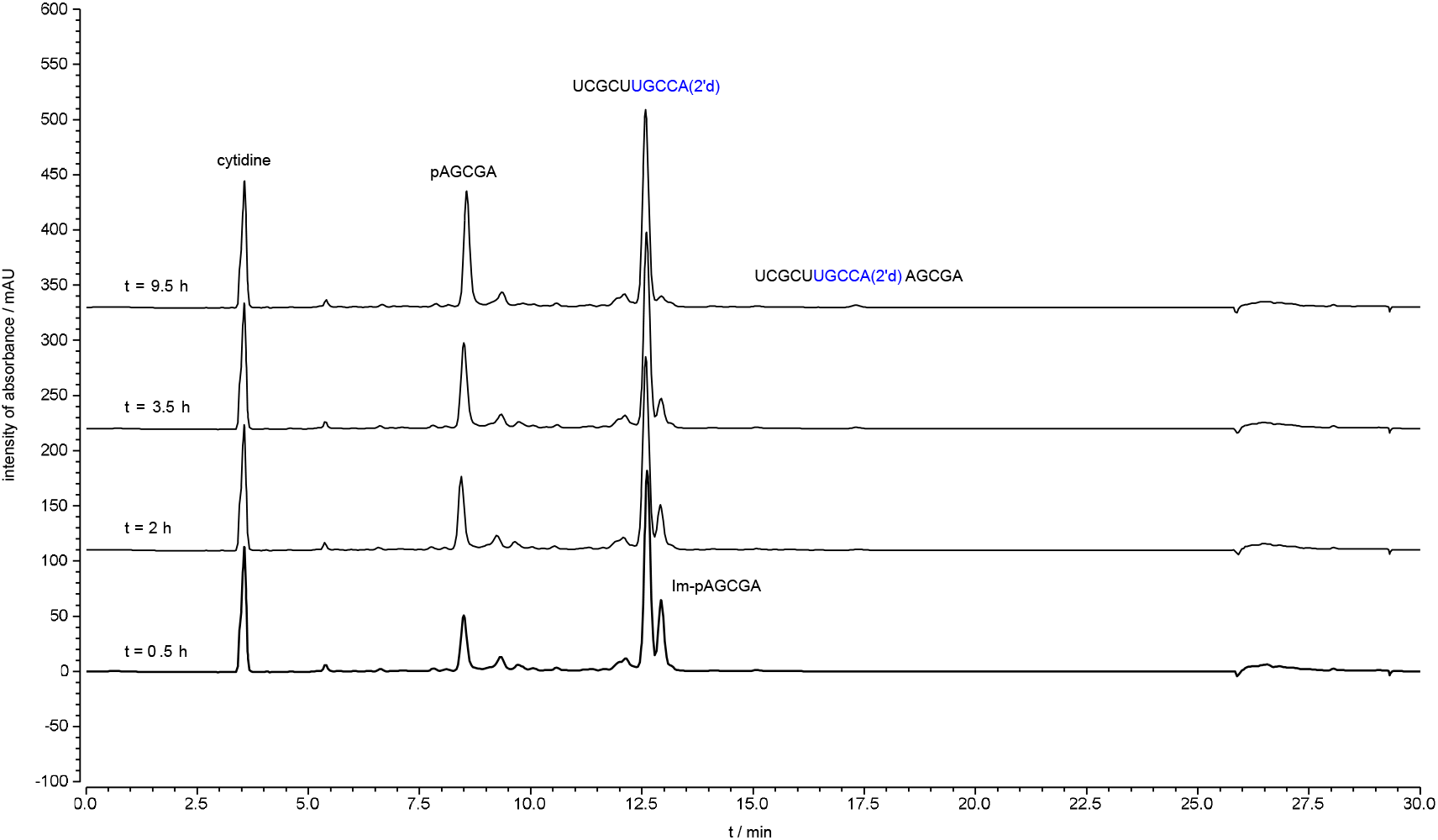
Stacked HPLC traces of loop-closing ligation with UGCCA(2’d) overhang. Loop duplex sequence: `3’AGCGAp-Im` `5’UCGCUUGCCA(2’d)` Loop-closing ligation was monitored by HPLC with 260 nm UV detection. The solution was incubated at 20 °C and aliquots of 8 μL were injected into an HPLC at different time points. Peaks for the phosphate donor, phosphate acceptor strands and the product of loop-closing ligation are indicated. Conditions: 50 μL of reaction mixture, containing the phosphate donor strand (including Im-p-AGCGA and p-AGCGA, in total 50 μM), the phosphate acceptor strand (5’-UCGCUUGCCA(2’d)-3’, 50 μM), cytidine (internal standard, 200 μM), NaCl (200 mM), MgCl_2_ (50 mM), *N*-MeIm (50 mM) and HEPES (50 mM, pH 8), was incubated at 20 °C.

**Figure S17.**
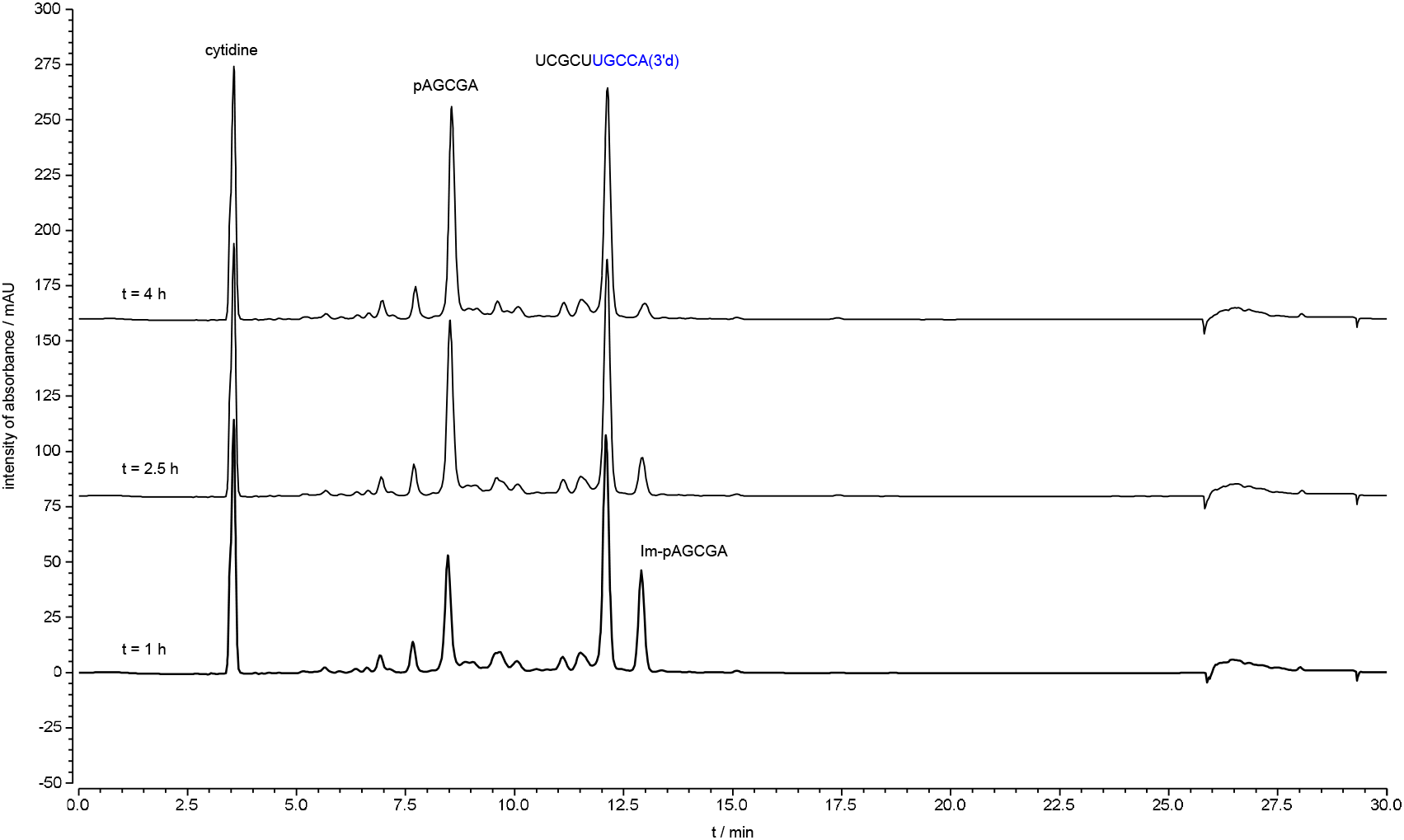
Stacked HPLC traces of loop-closing ligation with UGCCA(3’d) overhang. Loop duplex sequence: `3’AGCGAp-Im` `5’UCGCUUGCCA(3’d)` Loop-closing ligation was monitored by HPLC with 260 nm UV detection. The solution was incubated at 20 °C and aliquots of 8 μL were injected into an HPLC at different time points. Peaks for the phosphate donor, phosphate acceptor strands and the product of loop-closing ligation are indicated. Conditions: 50 μL of reaction mixture, containing the phosphate donor strand (including Im-p-AGCGA and p-AGCGA, in total 50 μM), the phosphate acceptor strand (5’-UCGCUUGCCA(3’d)-3’, 50 μM), cytidine (internal standard, 200 μM), NaCl (200 mM), MgCl_2_ (50 mM), *N*-MeIm (50 mM) and HEPES (50 mM, pH 8), was incubated at 20 °C.

**Figure S18.**
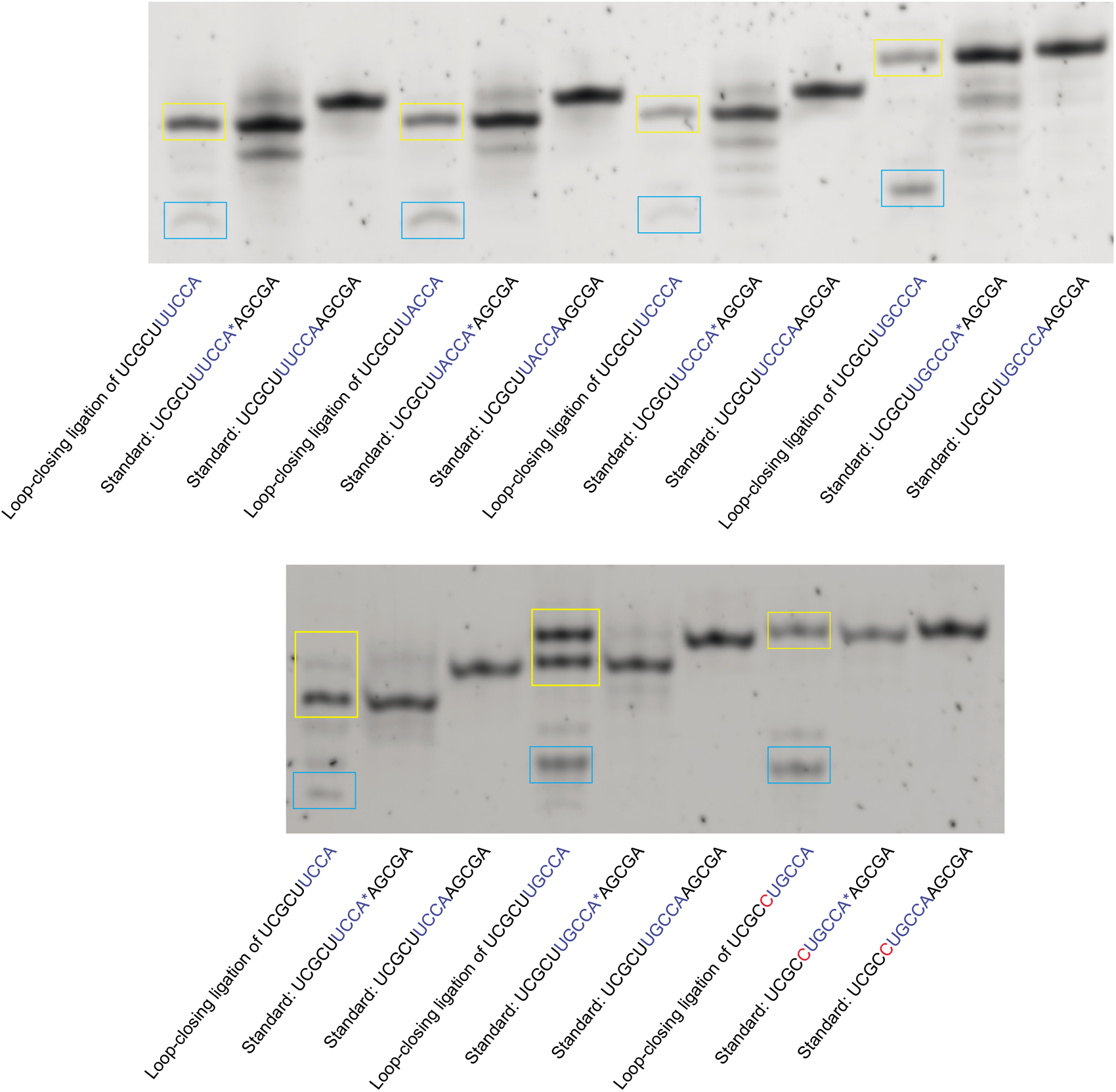
Characterisation of the regioselectivity of loop-closing ligations by PAGE. Starting materials (9 mer to 11 mer, Table 1) are indicated in blue boxes. Loop-closing ligation products are indicated in yellow boxes. A synthesised all-3’,5’-linkage authentic standard and an authentic standard with one 2’,5’-linkage at the loop-closing position were both run in parallel on the gel for comparison. A*A indicated the 2’,5’-linkage between these two nucleosides. The gel was stained by using SYBR Gold Nucleic Acid Gel Stain (no dye-labelling of those oligos) before imaging. The newly formed phosphodiester bond was predominantly 2’,5’-linked for those tested.

**Figure S19.**
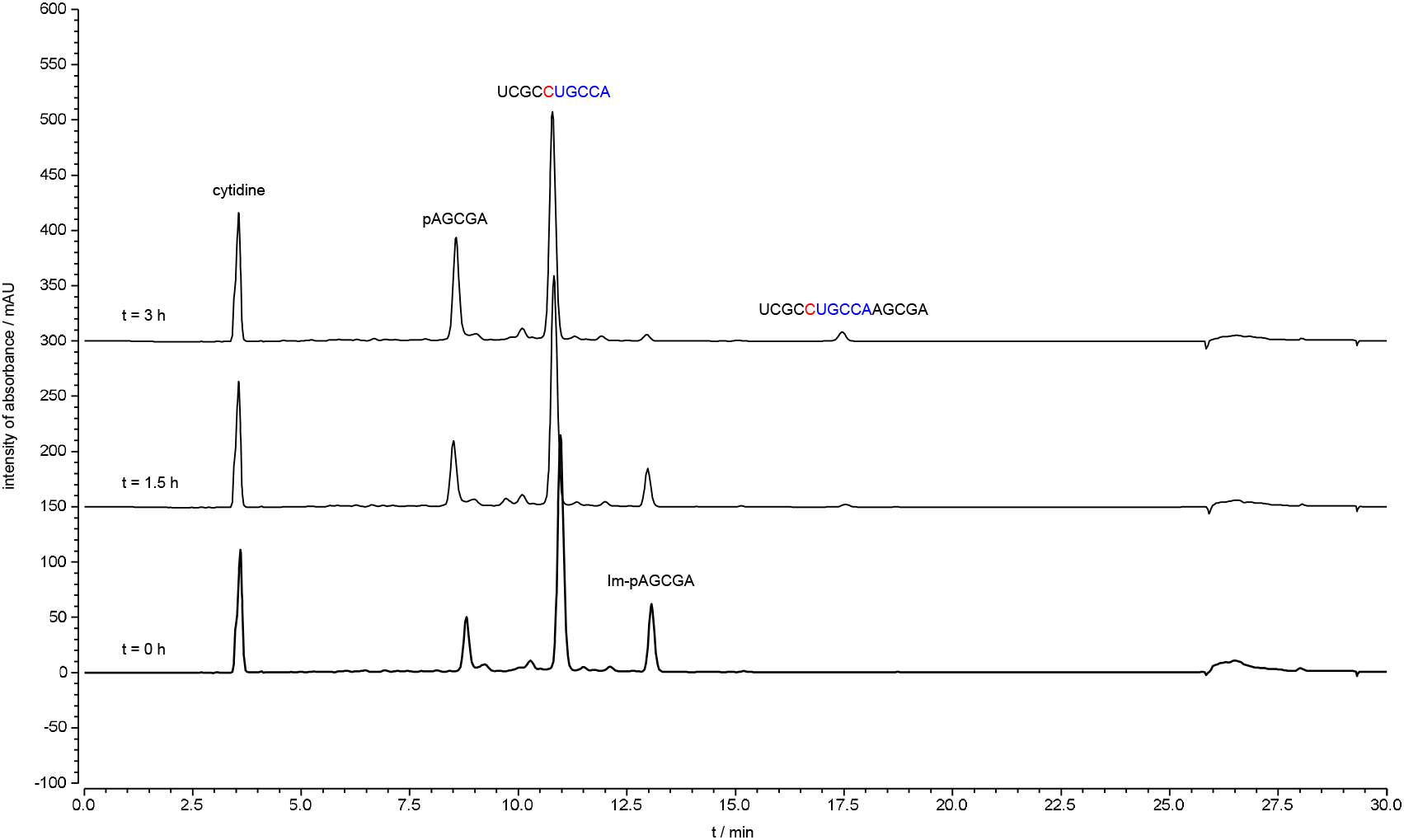
Stacked HPLC traces of loop-closing ligation with CUGCCA overhang on 3’-end and A on 5’-end. Loop duplex sequence: `3’AGCGAp-Im` 5’UCGCCUGCCA Loop-closing ligation was monitored by HPLC with 260 nm UV detection. The solution was incubated at 20 °C and aliquots of 8 μL were injected into an HPLC at different time points. Peaks for the phosphate donor, phosphate acceptor strands and the product of loop-closing ligation are indicated. Conditions: 50 μL of reaction mixture, containing the phosphate donor strand (including Im-p-AGCGA and p-AGCGA, in total 50 μM), the phosphate acceptor strand (5’-UCGCCUGCCA-3’, 50 μM), cytidine (internal standard, 200 μM), NaCl (200 mM), MgCl_2_ (50 mM), *N*-MeIm (50 mM) and HEPES (50 mM, pH 8), was incubated at 20 °C.

**Figure S20.**
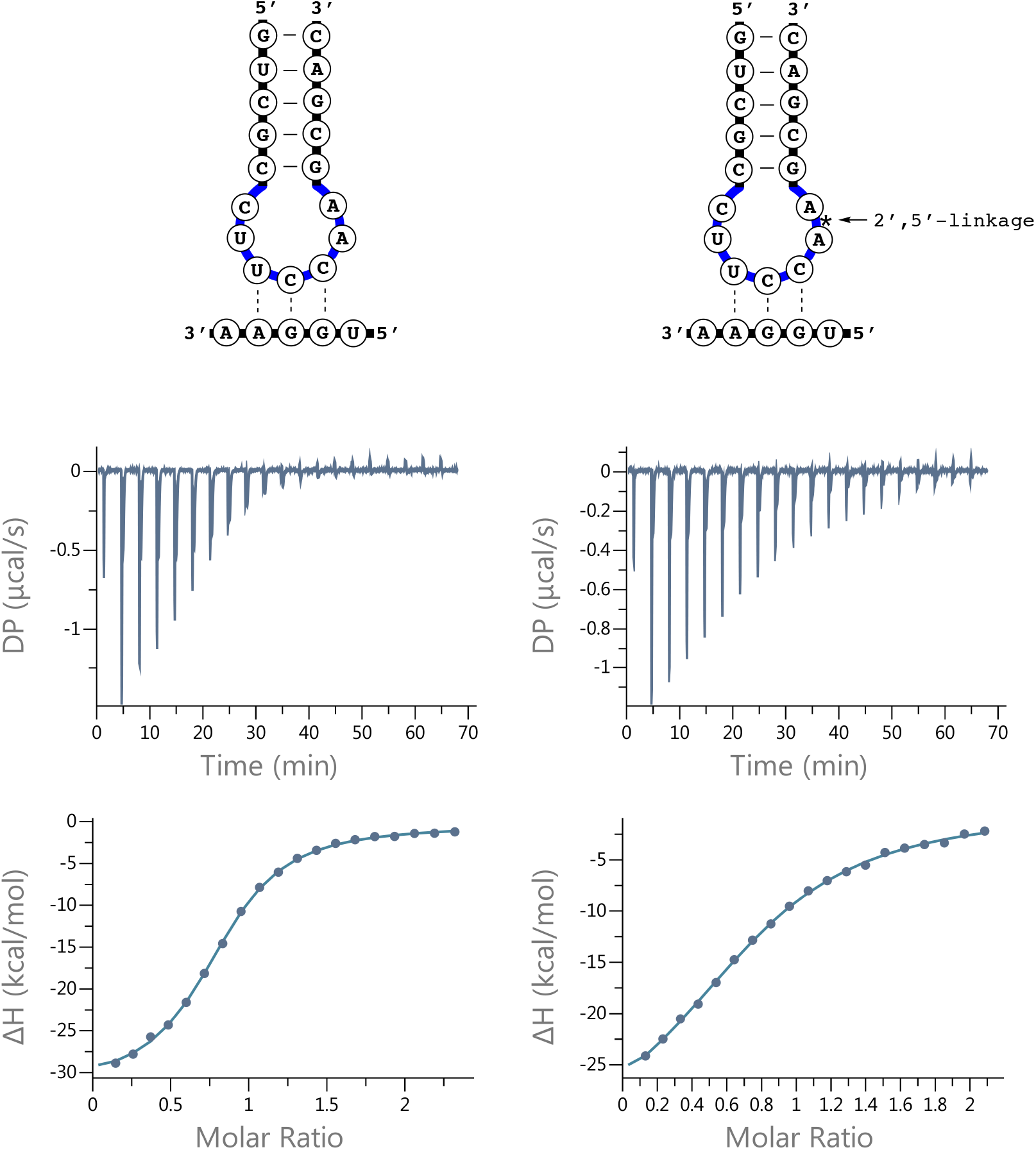
ITC data for binding of anticodon loop to a penta-nucleotide containing the corresponding codon. Left: using a loop containing all 3’,5’ linkages. Right: using a loop containing a 2’,5’ linkage between the positions equivalent to tRNA base 37 and 38.

**Figure S21.**
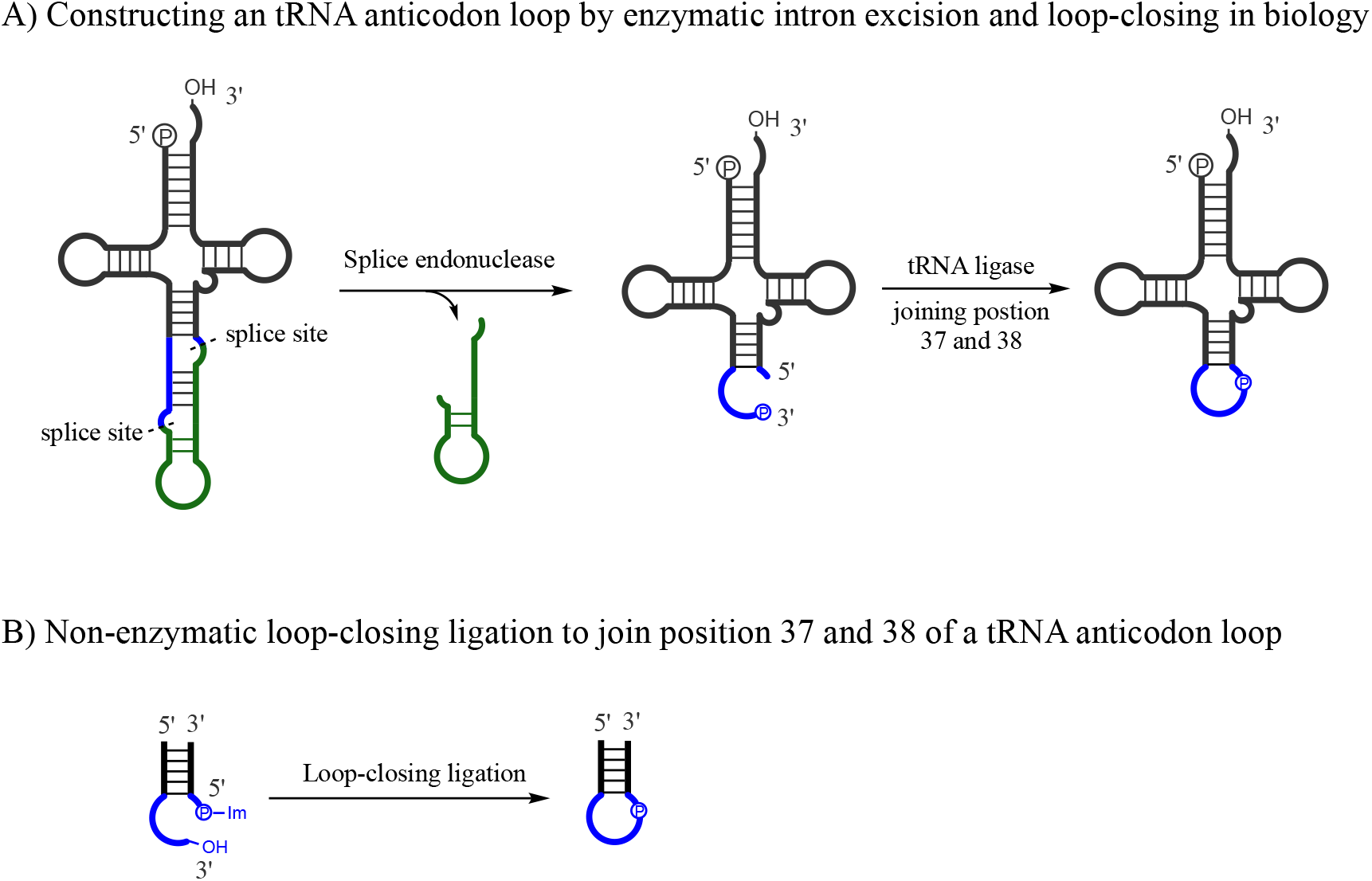
Constructing an anticodon loop structure by loop-closing ligation reminiscent of the pre-tRNA processing of the anticodon loop in biology.

**Figure S22.**
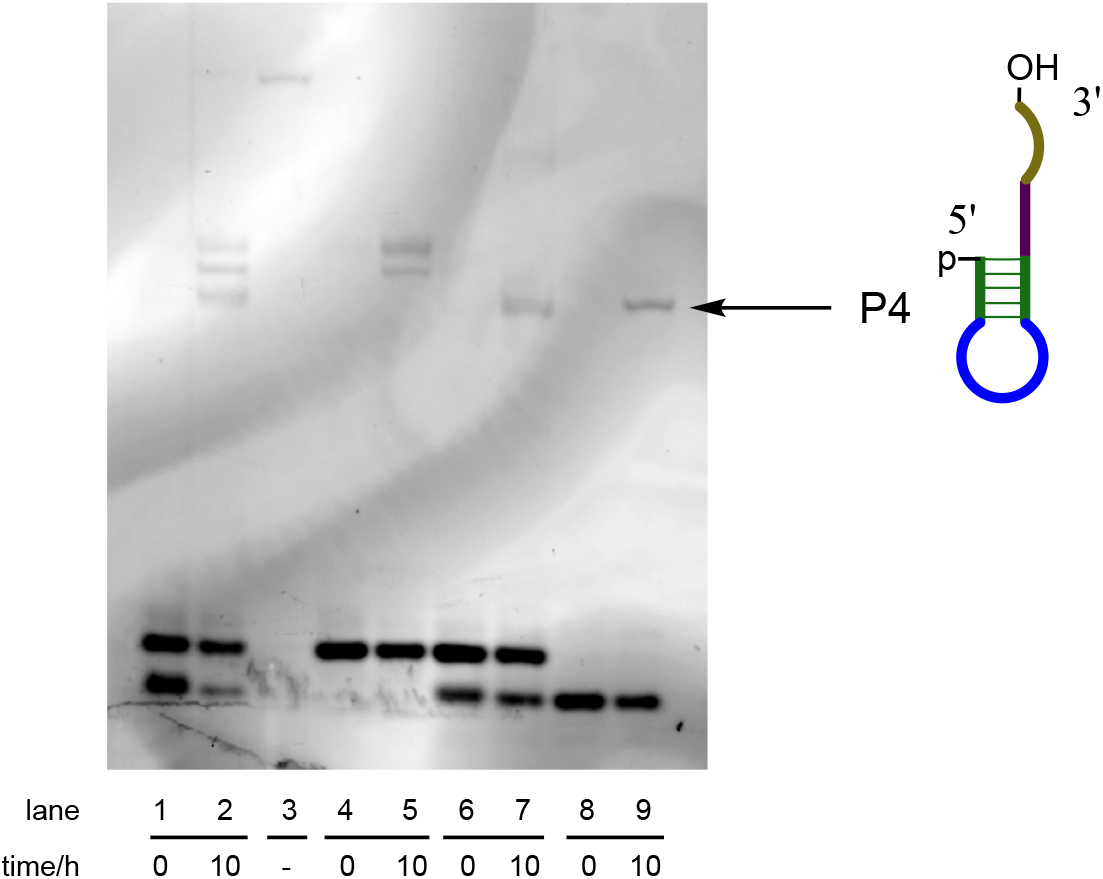
Direct assembly a minihelix RNA structure by loop-closing ligation (SYBR gold staining of the RNA gel shown in Figure 2). Lane 1&2, assembly reaction of Frg-1, Im-p-Frg-2 and Im-p-Frg-3; Lane 3, authentic standard of the minihelix RNA; Lane 4&5, reaction of Frg-1 and Im-p-Frg-3; Lane 6&7, reaction of Frg-1, Im-p-Frg-2 and p-Frg-3; Lane 8&9, reaction of p-Frg-2 and Im-p-Frg-3.

**Figure S23.**
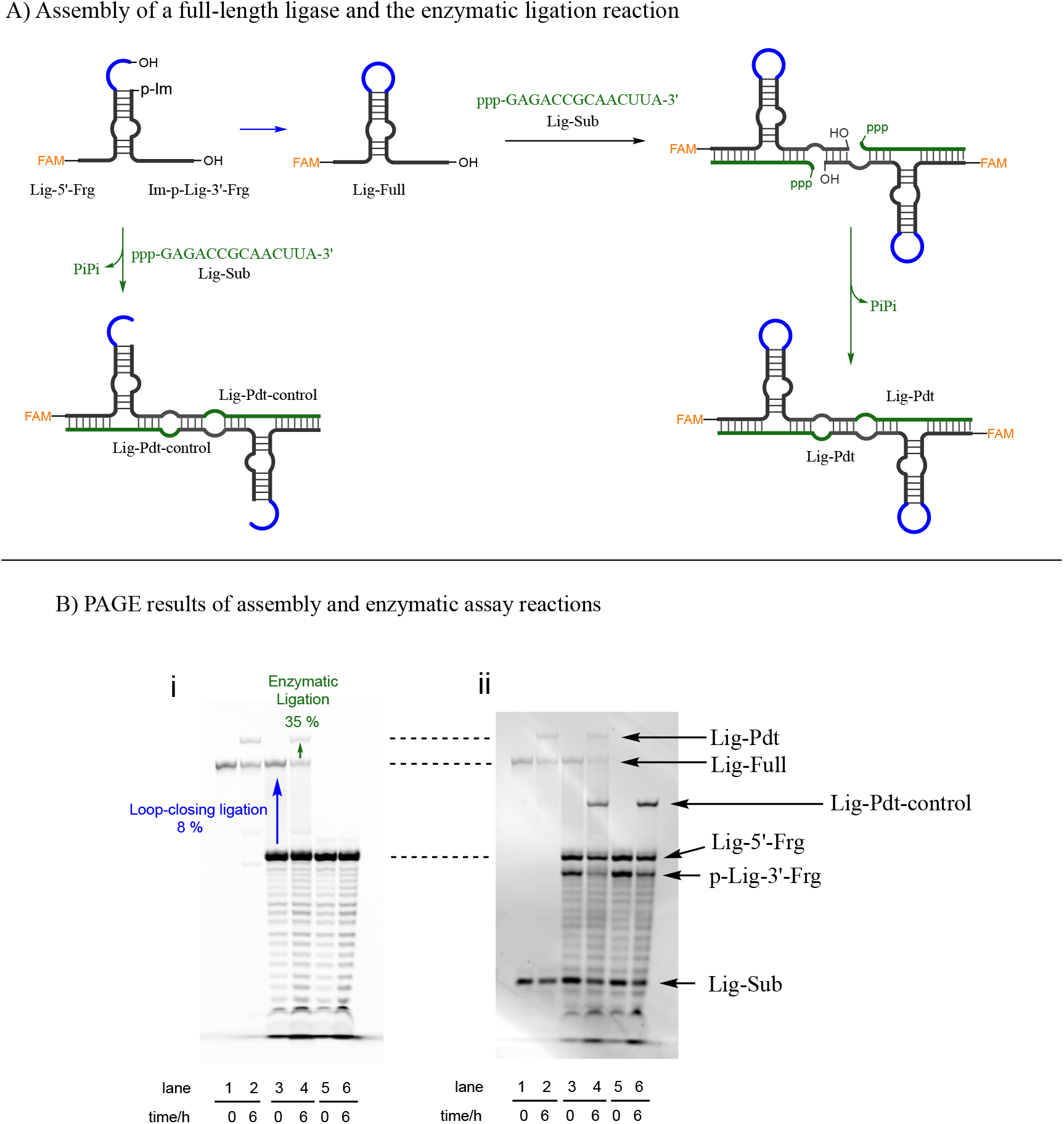
Direct assembly of the Joyce ligase ribozyme and the enzymatic ligation assay. A) Reaction scheme of the assembly of the ribozyme ligase and its subsequent enzymatic reaction. B) Representative PAGE gel electrophoresis for the assembly reaction and the enzymatic assay. i) Imaging based on FAM-labelling; ii) Imaging the same gel after SYBR-gold staining. Lane 1-2, positive control reaction of the Joyce ligase by using a pre-synthesised full-length ribozyme; Lane 3-4, enzymatic ligation reaction after loop-closing ligation; Lane 5-6, negative control without preceding loop-closing ligation.

**Table S1.** The reaction rates of loop-closing ligation depend on the concentration of N-methylimidazole (N-MeIm). The blue colour highlights the reference condition. Corrected yield = Observed yield divided by the initial fraction of Im-p-AGCGA present in the pre-synthesized mixture of Im-p-AGCGA & p-AGCGA (for synthetic methods see the SI). Reaction half-life, *t*_1/2_, is the combined rate of first-order consumption of Im-p-AGCGA resulting from both loop-closing ligation and the competing hydrolysis. All yields and half-lives are average values from at least two independent experiments.

| Phosphate donor | Phosphate acceptor sequence |  | pH | Temperature | NaCl (mM) | MgCl<sub>2</sub> (mM) | *N*-MeIm (mM) | Observed yield | Corrected yield | Reaction half-life (h) |
| --- | --- | --- | --- | --- | --- | --- | --- | --- | --- | --- |
|  | Stem | Overhang |  |  |  |  |  |  |  |  |
| Im-p-AGCGA | UCGCU | UGCCA | 8.0 | 20 °C | 200 | 50 | 0 | 9 % | 16 % | 40 |
|  |  |  | 8.0 | 20 °C | 200 | 50 | 10 | 18 % | 30 % | 7.5 |
|  |  |  | 8.0 | 20 °C | 200 | 50 | 20 | 17 % | 28 % | 3.3 |
|  |  |  | 8.0 | 20 °C | 200 | 50 | 50 | 18 % | 30 % | 1.8 |
|  |  |  | 8.0 | 20 °C | 200 | 50 | 80 | 18 % | 30 % | 1.3 |
|  |  |  | 8.0 | 20 °C | 200 | 50 | 100 | 18 % | 30 % | 1.0 |
|  |  |  | 8.0 | 20 °C | 200 | 50 | 200 | 17 % | 28 % | 0.5 |
|  |  |  | 8.0 | 30 °C | 200 | 50 | 50 | 16 % | 27 % | -- |
|  |  |  | 8.0 | 4 °C | 200 | 50 | 50 | 27 % | 45 % | -- |

**Table S2.**
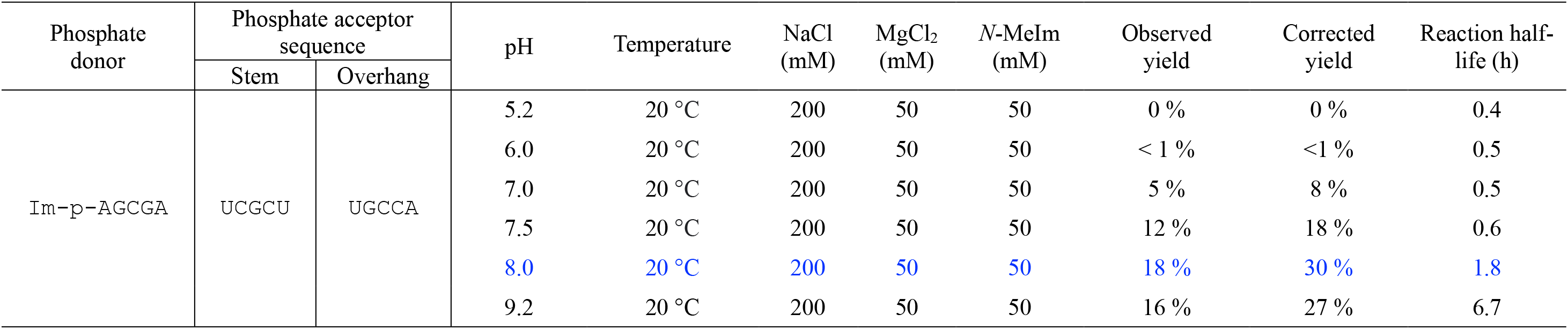
The pH-dependence of the loop-closing ligation. The blue colour highlights the reference condition. All yields and half-lives are average values from at least two independent experiments.

**Table S3.** The yields of loop-closing ligation depend on concentration of MgCl_2_. The blue colour highlights the reference condition. All yields and half-lives are average values from at least two independent experiments.

| Phosphate donor | Phosphate acceptor sequence |  | pH | Temperature | NaCl (mM) | MgCl<sub>2</sub> (mM) | *N*-MeIm (mM) | Observed yield | Corrected yield | Reaction half-life (h) |
| --- | --- | --- | --- | --- | --- | --- | --- | --- | --- | --- |
|  | Stem | Overhang |  |  |  |  |  |  |  |  |
| Im-p-AGCGA | UCGCU | UGCCA | 8.0 | 20 °C | 200 | 0 | 50 | 2 % | 3 % | 0.8 |
|  |  |  | 8.0 | 20 °C | 200 | 10 | 50 | 6 % | 10 % | 1.1 |
|  |  |  | 8.0 | 20 °C | 200 | 20 | 50 | 13 % | 21 % | 1.4 |
|  |  |  | 8.0 | 20 °C | 200 | 50 | 50 | 19 % | 31 % | 1.8 |
|  |  |  | 8.0 | 20 °C | 200 | 100 | 50 | 25 % | 41 % | 2.4 |
|  |  |  | 8.0 | 20 °C | 200 | 200 | 50 | 29 % | 48 % | 2.9 |
|  |  |  | 8.0 | 20 °C | 200 | 500 | 50 | 29 % | 48 % | 4.9 |

**Table S4.** The yields of loop-closing ligation depend on concentration of NaCl. The blue colour highlights the reference condition. All yields and half-lives are average values from at least two independent experiments.

| Phosphate Donor | Phosphate acceptor sequence |  | pH | Temperature | NaCl (mM) | MgCl<sub>2</sub> (mM) | *N*-MeIm (mM) | Observed yield | Corrected yield | Reaction half-life (h) |
| --- | --- | --- | --- | --- | --- | --- | --- | --- | --- | --- |
|  | Stem | Overhang |  |  |  |  |  |  |  |  |
| Im-p-AGCGA | UCGCU | UGCCA | 8.0 | 20 °C | 0 | 0 | 50 | 1 % | 2 % | 0.4 |
|  |  |  | 8.0 | 20 °C | 100 | 0 | 50 | 3 % | 4 % | 0.7 |
|  |  |  | 8.0 | 20 °C | 200 | 0 | 50 | 4 % | 6 % | 1.0 |
|  |  |  | 8.0 | 20 °C | 500 | 0 | 50 | 5 % | 10 % | 1.5 |
|  |  |  | 8.0 | 20 °C | 1000 | 0 | 50 | 6 % | 15 % | 2.2 |
|  |  |  | 8.0 | 20 °C | 2000 | 0 | 50 | 16 % | 22 % | 3.9 |

## References

1. J. R. Wyatt, I. Jr. Tinoco, RNA structural elements and RNA function. In the RNA World, (ed. Gesteland, R. F., Atkins, J. F.), pp. 465–97. (Cold Spring Harbor Lab. Press, 1993).

2. J. A. Doudna, Tertiary motifs in RNA structure and folding. Angew. Chem. Int. Ed. 38, 2326–2343 (1999).

3. E. A. Doherty, J. A. Doudna, Ribozyme structures and mechanisms. Annu. Rev. Biophys. Biomol. Struct., 30, 457–75 (2001).

4. R. T. Batey, R. P. Rambo, M. J. Fedor, J. R. Williamson, The catalytic diversity of RNAs. Nat. Rev. Mol. Cell Biol.6, 399–412 (2005).

5. S. E. Butcher, A. M. Pyle, The molecular interactions that stabilize RNA tertiary structure: RNA motifs, patterns, and networks. Acc. Chem. Res. 44, 12, 1302–1311 (2011).

6. J. P. Ferris, Montmorillonite-catalysed formation of RNA oligomers: the possible role of catalysis in the origins of life. Phil. Trans. R. Soc. B, 361, 1777–1786 (2006)

7. L. E. Orgel, Molecular replication. Nature 358, 203–209 (1992).

8. J. W. Szostak, The eightfold path to non-enzymatic RNA replication. J. Syst. Chem. 2012, 3, 2 (2012)

9. F. Wachowius, P. Holliger, Non-enzymatic assembly of a minimized RNA polymerase ribozyme. ChemSystemsChem. 1, No. e1900004 (2019).

10. L. Zhou, D. K. O’Flaherty, J. W. Szostak, Assembly of a ribozyme ligase from short oligomers by nonenzymatic ligation. J. Am. Chem. Soc. 142, 15961−5965 (2020).

11. J. A. Doudna, S. Couture, J. W. Szostak, A multisubunit ribozyme that is a catalyst of and template for complementary strand RNA synthesis. Science 251, 1605–1608 (1991).

12. T. A. Rogers, G. E. Andrews, L. Jaeger, W. W. Grabow, Fluorescent monitoring of RNA assembly and processing using the split-spinach aptamer. ACS Synth. Biol. 4, 162–166 (2015).

13. A. Akoopie, U. F. Müller, Lower temperature optimum of a smaller, fragmented triphosphorylation ribozyme. Phys. Chem. Chem. Phys. 18, 20118–20125 (2016).

14. K. F. Tjhung, M. N. Shokhirev, D. P. Horning, G. F. Joyce, An RNA polymerase ribozyme that synthesizes its own ancestor. Proc. Natl. Acad. Sci. USA 117, 2906–2913 (2020).

15. R. Naylor, P. T. Gilham, Studies on some interactions and reactions of oligonucleotides in aqueous solution. Biochemistry 5, 2722–2728 (1966).

16. F. Diederich, P. J. Stang, Eds., Templated Organic Synthesis Wiley-VCH, Weinheim, 2000.

17. D. Herschlag, RNA chaperones and the RNA folding problem. J. Biol. Chem. 270, 20871–20874 (1995).

18. L. Rajkowitsch, D. Chen, S. Stampfl, K. Semrad, C. Waldsich, O. Mayer, M. F. Jantsch, R. Konrat, U. Bläsi, R. Schroeder, RNA chaperones, RNA annealers and RNA helicases. RNA Biol. 4, 118–30 (2007).

19. O.C. Uhlenbeck, Keeping RNA happy. RNA 1, 4–6 (1995).

20. L.-F. Wu, M. Su, Z. Liu, S. Bjork, J. D. Sutherland, Interstrand aminoacyl-transfer in a tRNA acceptor stem mimic. J. Am. Chem. Soc. 143, 11836–11842 (2021).

21. T. Walton, W. Zhang, L. Li, C. P. Tam, J. W. Szostak, The mechanism of nonenzymatic template copying with imidazole-activated nucleotides. Angew. Chem. Int. Ed. 58, 10812–10819 (2019).

22. A. Mariani, D. A. Russell, T., Javelle, J. D. Sutherland, A light-releasable potentially prebiotic nucleotide activating agent. J. Am. Chem. Soc. 140, 8657–8661 (2016).

23. K. Tamura, P. Schimmel, Chiral-selective aminoacylation of an RNA minihelix. Science 305, 1253 (2004).

24. E. V. Puglisi, J. D. Puglisi, J. R. Williamson, U. L. RajBhandary, NMR analysis of tRNA acceptor stem microhelices: Discriminator base affects tRNA conformation at the 3′-end. Proc. Natl. Acad. Sci. USA 91, 11467–11471 (1994).

25. R. Rohatgi, D. P. Bartel, J. W. Szostak, Nonenzymatic, template-directed ligation of oligoribonucleotides is highly regioselective for the formation of 3′-5′ phosphodiester bonds. J. Am. Chem. Soc. 118, 3340–3344 (1996).

26. A. Kanavarioti, C. F. Bernasconi, D. L. Doodokyan, D. J. Alberas, Magnesium ion catalyzed phosphorus-nitrogen bond hydrolysis in imidazolide-activated nucleotides. Relevance to template-directed synthesis of polynucleotides J. Am. Chem. Soc. 111, 18, 7247–7257 (1989).

27. G. Varani, Exceptionally stable nucleic acid hairpins. Annu. Rev. Biophys. Biomol. Struct. 24, 379–404 (1995).

28. J. Wolters, The nature of preferred hairpin structures in 16S-like rRNA variable regions. Nucleic Acids Res. 20, 1843–1850 (1992).

29. P. C. Bevilacqua, J. M. Blose, Structures, kinetics, thermodynamics, and biological functions of RNA hairpins. Annu. Rev. Phys. Chem. 59, 79–103 (2008).

30. P. Svoboda, A. Di Cara, Hairpin RNA: a secondary structure of primary importance. Cell. Mol. Life Sci. 63, 901–918 (2006).

31. A. E. Engelhart, M. W. Powner, J. W. Szostak, functional RNAs exhibit tolerance for non-heritable 2′,5′ versus 3′,5′ backbone heterogeneity. Nat. Chem. 5, 390–394 (2013).

32. O. C. Uhlenbeck, Complementary oligonucleotide binding to transfer RNA. J. Mol. Biol. 65, 25–41 (1972).

33. J. Popow, A. Scheiffer, J. Martinez, Diversity and roles of (t)RNA ligases. Cell. Mol. Life Sci. 69, 2657–2670 (2012).

34. C. Francklyn, P. Schimmel, Amnoacylation of RNA minihelices with alanine. Proc. Natl. Acad. Sci. USA 337, 478–481 (1989).

35. H. W., Pley, K. M. Flaherty, D. B. McKay, Three-dimensional structure of a hammerhead ribozyme. Nature 372, 68–74 (1994).

36. M. P. Robertson, G. F. Joyce, Highly efficient self-replicating RNA enzymes. Chem. Bio. 21, 238–245 (2014).

37. J. H. Davis, J. W. Szostak, Isolation of high-affinity GTP aptamers from partially structured RNA libraries. Proc. Natl. Acad. Sci. USA 99, 11616–11621 (2002).

38. F. Chizzolini, L. F. M. Passalacqua, M. Oumais, A. Dingilian, J. W. Szostak, A. Luptak, Large phenotypic enhancement of structured random RNA pools. J. Am. Chem. Soc.142, 1941–1951 (2020).

39. M. Chastain, I. Jr. Tinoco, Structural elements in RNA. Prog. Nucleic Acid Res. Mol. Biol. 41, 131–177 (1991).

40. P. B. Moore, Structural motifs in RNA. Annu. Rev. Biochem. 68, 287–300 (1999).

41. D. K. Hendrix, S. E. Brenner, S. R. Holbrook, RNA structural motifs: building blocks of a modular biomolecule. Q. Rev. Biophys. 38, 221–243 (2005).

42. Z. Liu, L.-F. Wu, J. Xu, C. Bonfio, D. A. Russell, J. D. Sutherland, Harnessing chemical energy for the activation and joining of prebiotic building blocks. Nat. Chem. 12, 1023–1028 (2020).

43. S. J. Zhang, D. Duzdevich, J. W. Szostak, Potentially prebiotic activation chemistry compatible with nonenzymatic RNA copying. J. Am. Chem. Soc. 142, 14810–14813 (2020).

44. A. Kanai, Disrupted tRNA genes and tRNA fragments: a perspective on tRNA gene evolution. Life 5, 321–331 (2015).

45. M. W. Gray, V. Gopalan, Piece by piece: building a ribozyme. J. Biol. Chem. 295, 2313–2323 (2020).

